# Genetic Screen Uncovers a Dual Role for Phospholipids in Mitochondrial-Derived Compartment Biogenesis

**DOI:** 10.1101/2023.02.16.528844

**Authors:** Tianyao Xiao, Alyssa M. English, J. Alan Maschek, James E. Cox, Adam L. Hughes

**Affiliations:** Department of Biochemistry, University of Utah School of Medicine, Salt Lake City, UT, 84112, USA; Metabolomics Core Research Facility, University of Utah, Salt Lake City, UT, 84112, USA; Department of Nutrition & Integration Physiology, University of Utah College of Health, Salt Lake City, UT, 84112, USA

**Author notes:** Correspondence: Department of Biochemistry, University of Utah School of Medicine, 15 N. Medical Drive East, RM 4100, Salt Lake City, UT, 84112.

## Abstract

Cells utilize multiple mechanisms to maintain mitochondrial homeostasis in response to stress. We recently identified a cellular structure, called the mitochondria-derived compartment (MDC), that is generated from mitochondria in response to amino acid overabundance. MDCs selectively sequester proteins from mitochondria for subsequent degradation, and loss of MDCs sensitizes cells to amino acid stress. Here, we conducted a microscopy-based screen in budding yeast to identify factors that regulate MDC formation. We found that levels of two mitochondrial phospholipids, cardiolipin (CL) and phosphatidylethanolamine (PE), regulate MDC biogenesis in opposing directions. CL depletion impairs MDC biogenesis, whereas PE reduction leads to constitutive MDC formation. Additionally, in response to MDC-inducing agents, cellular and mitochondrial PE declines in an amino acid-dependent manner. Overexpressing mitochondrial PE synthesis pathway components suppresses MDC biogenesis during amino acid stress. Altogether, our data indicate a requirement for CL in MDC biogenesis, and suggest that PE depletion may serve as a regulatory signal for MDC formation downstream of MDC-inducing metabolic stress.

**SUMMARY:** Xiao et al. identify distinct roles for the phospholipids cardiolipin and phosphatidylethanolamine in mitochondria-derived compartment (MDC) biogenesis. They show that cardiolipin is required for MDC formation, whereas a decline in phosphatidylethanolamine in response to amino acid stress triggers MDC biogenesis.

## INTRODUCTION

Mitochondria are hubs for cellular metabolism (Friedman & Nunnari, 2014; Spinelli & Haigis, 2018). Disrupted mitochondrial homeostasis contributes to aging and numerous metabolic diseases (Wallace, 2005; Lin and Beal, 2006; Kudryavtseva et al., 2016). To maintain mitochondrial function under pathological stress, cells utilize various quality control and signaling pathways, including mitochondrial proteases (Quirós et al., 2015), the ubiquitin-proteasome system (Bragoszewski et al., 2017; Ravanelli et al., 2020), mitophagy (Palikaras et al., 2018; Pickles et al., 2018), mitochondrial-derived vesicles (Sugiura et al., 2014; Picca et al., 2020; König et al., 2021), mitochondrial fission and fusion (Giacomello et al., 2020), the mitochondrial unfolded protein response (Melber and Haynes, 2018) and multiple mitoprotein-induced stress responses (Song et al., 2021; Boos et al., 2020). In recent studies, our group identified a new mechanism by which cells regulate mitochondrial homeostasis—through formation of <u>m</u>itochondrial-<u>d</u>erived <u>c</u>ompartments (MDCs) (Hughes et al., 2016; English et al., 2020; Schuler et al., 2021). In *Saccharomyces cerevisiae*, MDCs are dynamic mitochondrial-associated structures generated from mitochondrial membranes that appear to contain distinct lumens (Schuler et al., 2021; English et al., 2020). MDC biogenesis is induced by elevated intracellular amino acid abundance (Schuler et al., 2021), which is a cellular stress that occurs during the progression of aging, metabolic disorders, and neurodegenerative diseases (Wellen and Thompson, 2010; Hughes and Gottschling, 2012; Aliu et al., 2018; Hughes et al., 2020; Ruiz et al., 2021). In previous studies, we identified several perturbations that trigger MDC formation by causing a build-up of intracellular amino acids, including impairing vacuolar amino acid storage using the vacuolar H^+^-ATPase inhibitor concanamycin A (ConcA), and preventing amino acid incorporation into proteins using rapamycin (Rap) or cycloheximide (CHX), which inhibit translation (Schuler et al., 2021; English et al., 2020). The induction of MDCs by ConcA, Rap, and CHX is suppressed by depleting amino acids from the media (Schuler et al., 2021).

Under amino acid stress, MDCs selectively sequester Tom70, a receptor of the mitochondrial outer membrane translocase (TOM) complex (Söllner et al., 1990; Steger et al., 1990), as well as other mitochondrial outer membrane (OMM) proteins and mitochondrial carrier proteins, diminishing the levels of these proteins in mitochondria (Hughes et al., 2016; Schuler et al., 2021). In contrast, mitochondrial intermembrane space (IMS) proteins, matrix proteins, and the majority of mitochondrial inner membrane (IMM) proteins are excluded from MDCs (Hughes et al., 2016). After formation, MDCs are separated from the mitochondrial tubule by fission and delivered to the vacuole for degradation (Hughes et al., 2016). Thus, MDCs are cargo-selective structures that remodel the mitochondrial proteome in response to increased intracellular amino acid levels. Failure to form MDCs alters amino acid catabolism and sensitizes cells to amino acid stress (Schuler et al., 2021).

Using super-resolution microscopy, we previously showed that MDCs are closely associated with both mitochondria and the endoplasmic reticulum (ER) (English et al., 2020). MDC generation occurs at the contact sites between the ER and mitochondria, and requires proteins that localize to these sites, including the ER–mitochondria encounter structure (ERMES) complex and the ERMES-associated GTPase Gem1 (Kornmann et al., 2009, 2011; English et al., 2020). Various genetic suppressors that rescue defects in ERMES mutants differ in their ability to restore MDC formation (Tan et al., 2013; Lang et al., 2015; John Peter et al., 2017; English et al., 2020). As a result, it remains unclear how ER-mitochondria contacts regulate MDC biogenesis. Additionally, other factors involved in relaying signals and facilitating MDC formation remain unknown.

Here, we present results from a microscopy-based screen of the yeast non-essential deletion collection (Giaever et al., 2002) aimed at uncovering genes involved in MDC formation. As described below, we identified a number of gene deletions that enhance or suppress MDC formation, which led to discovery of dual roles for the non-bilayer forming phospholipids cardiolipin (CL) and phosphatidylethanolamine (PE) in MDC biogenesis.

## RESULTS

### An imaging-based screen identifies genetic regulators of MDC biogenesis

To identify genetic regulators of MDC biogenesis, we conducted a genome-wide microscopy-based screen using the yeast non-essential deletion collection (Giaever et al., 2002). We previously showed that MDCs are enriched with OMM proteins, including Tom70, but exclude most IMM and matrix proteins, such as Tim50 (Yamamoto et al., 2002; Hughes et al., 2016) (Fig. 1 A). Thus, we used the Synthetic Genetic Array (SGA) technique (Tong et al., 2001) to construct a yeast non-essential deletion library in which individual mutants contain endogenously tagged Tom70-GFP and Tim50-mCherry (mCh). Using high-throughput imaging, we examined the formation of Tom70-GFP-positive Tim50-mCh-negative foci (MDCs) in ∼5,000 deletion strains after treatment with the MDC inducer Rap (Schuler et al., 2021) (Fig. 1 B). The percentage of cells with MDCs in each mutant was quantified (Table S1). We found that the majority of deletion strains (3744 out of 4757 ORFs that were successfully screened) robustly formed MDCs in 20%-60% of cells (Fig. 1 C) when treated with Rap in 96-well plates, which is generally lower than we previously observed when cells were cultured in tubes or flasks (Schuler et al., 2021; English et al., 2020). Gene deletions that led to MDC formation in less than 20% or greater than 60% of cells were categorized as having potentially “low” and “high” MDC formation rates, respectively. These genes were subjected to gene ontology analysis with additional manual curation and are listed in Fig. 1 D.

**Figure 1.**
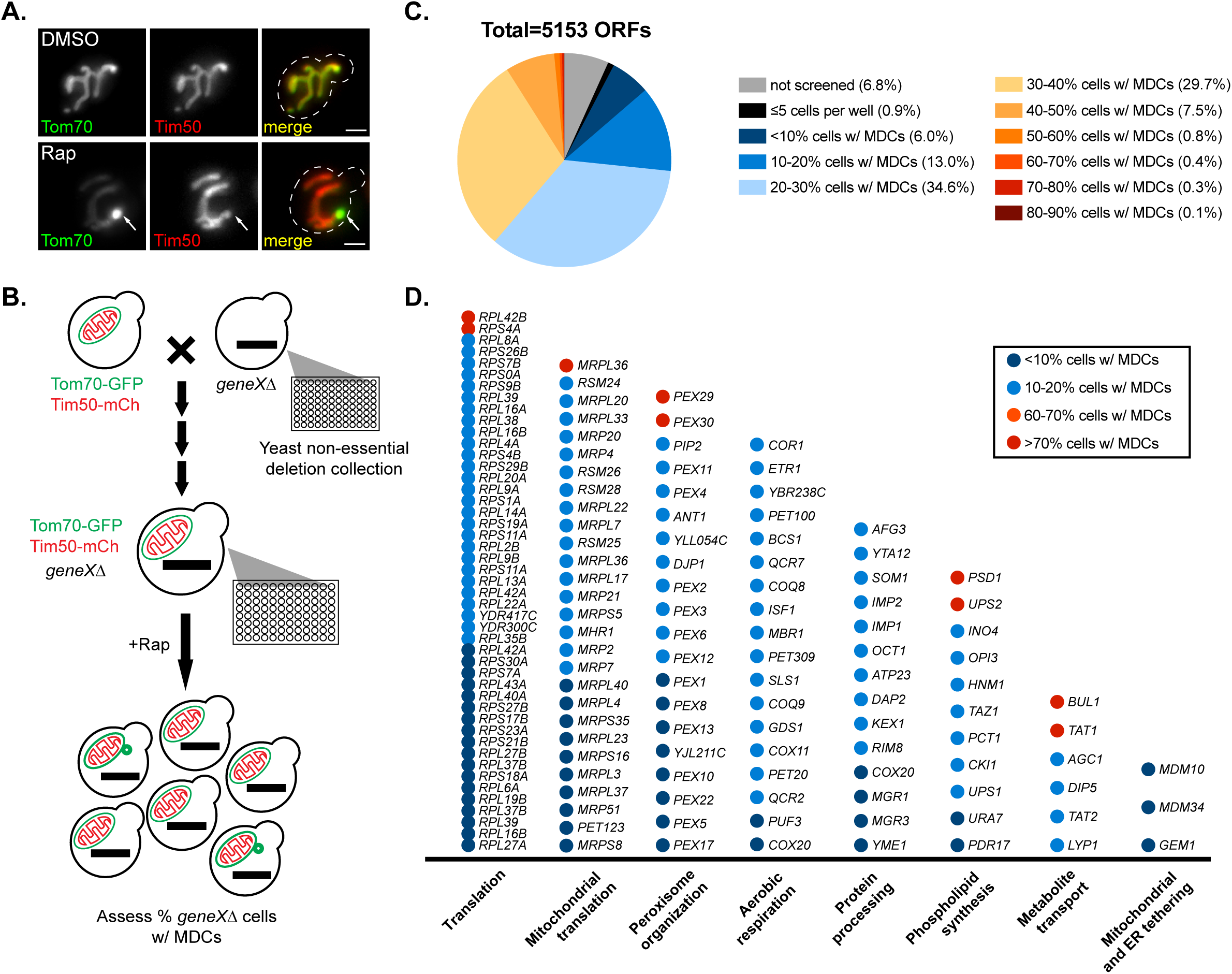
An imaging-based screen identifies genetic regulators of MDC biogenesis. **(A)** Widefield images of wild-type yeast cells endogenously expressing Tom70-yeGFP (Tom70-GFP) and Tim50-mCherry (Tim50-mCh) treated with DMSO or rapamycin (Rap) for 2 h. White arrows mark positions of MDCs. Scale bar = 2 µm. **(B)** Schematic of the genome-wide screen to identify genetic regulators of Rap-induced MDC biogenesis. A query strain containing endogenously expressing Tom70-GFP and Tim50-mCh was mated to the yeast non-essential deletion collection to obtain a new collection in which all mutants are labelled with Tom70-GFP and Tim50-mCh after several steps of selection. The collection was treated with Rap in 96-well plates for 2 h and imaged by automated microscopy. MDCs were identified as Tom70-GFP containing mitochondria-associated structures that are Tim50-mCh negative. Images were manually assessed and quantified to determine percentages of cells with MDCs in each well. **(C)** Fan plot of the MDC screen results showing ratio of mutants that could not screened (no growth or low image quality), grew poorly (≤ 5 cells per well), or formed MDCs in the indicated percentages of cells, to the total number of open reading frames (ORFs) contained in the yeast deletion collection. **(D)** Categories of gene deletions that led to decreased MDC biogenesis (≤20% cells form MDCs) or enhanced MDC biogenesis (≥60% cells form MDCs). For a complete list of all genes, their descriptions, and gene ontology analysis, see Table S1.

We successfully identified gene deletions that were previously shown to block MDC biogenesis, including ER-mitochondria tethering genes *MDM10*, *MDM34*, and *GEM1* (Kornmann et al., 2009, 2011; English et al., 2020) (Fig. 1 D). This indicated that the screening approach was capable of identifying genetic regulators of MDC formation. Through ontology analysis, we identified several gene categories that either positively or negatively regulate MDC biogenesis. These include a number of genes that are involved in cytoplasmic and mitochondrial translation, peroxisome organization, aerobic respiration, protein processing, phospholipid biosynthesis, and amino acid transport (Table S1, Fig. 1 D). We have not yet validated all of the hits from the screen. However, the large amount of gene deletions in each of the categories listed in Fig. 1 D suggests these biological processes likely impact MDC formation. Interestingly, consistent with previous observations that Rap triggers MDC induction by driving amino acid overabundance in cells (Schuler et al., 2021), deletion of several amino acid transporters decreased MDC formation rates, providing additional connections between MDCs and amino acid homeostasis.

### *UPS1* and *UPS2* regulate MDC biogenesis in an inverse manner

Among genes identified from the MDC screen, we found that *UPS1* and *UPS2*, which encode for IMS-localized intramitochondrial lipid transfer proteins of the PRELI family involved in CL and PE biosynthesis (Dee and Moffat, 2005; Sesaki et al., 2006; Tamura et al., 2009; Osman et al., 2009; Potting et al., 2010; Connerth et al., 2012; Tamura et al., 2012; Miyata et al., 2016), regulate MDC biogenesis in opposing directions (Fig. 1 D). CL and PE are the major non-bilayer forming phospholipids in cells (Osman et al., 2011). Both CL and PE play important roles in mitochondrial structure and function (Acoba et al., 2020; Basu Ball et al., 2018), and genetic depletion of both phospholipids in mitochondria causes synthetic lethality (Gohil et al., 2005). CL and PE are synthesized on the IMM using substrates obtained from the OMM. The movement of lipid precursors from the OMM to the IMM is mediated by Ups1 and Ups2 (Sesaki et al., 2006; Tamura et al., 2009; Osman et al., 2009; Potting et al., 2010; Connerth et al., 2012; Tamura et al., 2012; Miyata et al., 2016; Acoba et al., 2020). Specifically, Ups1 transports phosphatidic acid (PA) from the OMM to IMM for CL synthesis (Connerth et al., 2012), and Ups2 transports phosphatidylserine (PS) from the OMM to IMM for PE synthesis (Miyata et al., 2016; Aaltonen et al., 2016) (Fig. 2 A). Both Ups1 and Ups2 bind to Mdm35, which stabilizes them against proteolytic degradation (Miyata et al., 2016; Connerth et al., 2012; Potting et al., 2010). Deletion of *UPS1* blocks mitochondrial CL production (Tamura et al., 2009; Connerth et al., 2012). Similarly, loss of *UPS2* prevents PE synthesis in mitochondria (Miyata et al., 2016).

**Figure 2.**
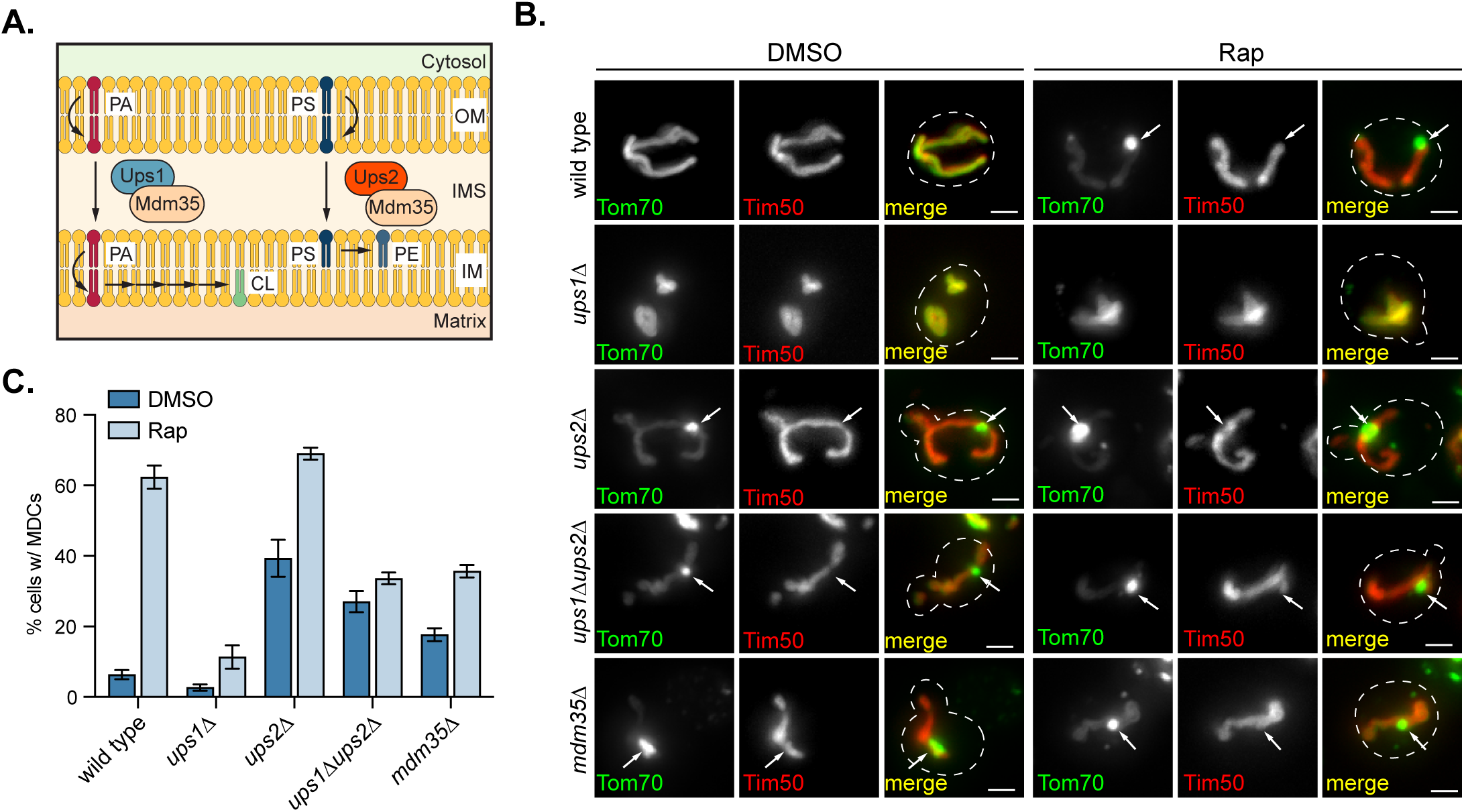
*UPS1* and *UPS2* regulate MDC biogenesis in opposing directions. **(A)** Model of intramitochondrial transport of substrates for CL and PE synthesis on the IMM mediated by IMS proteins Ups1 and Ups2. Ups1 transports PA for CL synthesis, and Ups2 translocates PS for PE synthesis. Ups1 and Ups2 are stabilized against degradation by binding to Mdm35. **(B)** Widefield images of wild-type cells or the indicated mutant yeast endogenously expressing Tom70-GFP and Tim50-mCh treated with DMSO or Rap for 2 h. White arrows mark positions of MDCs. Scale bar = 2 µm. **(C)** Quantification of (B) showing the percentage of cells with MDCs. N > 100 cells per replicate, error bars = SEM of three replicates.

From the initial screen, we found that loss of *UPS1* inhibited MDC formation, whereas deletion of *UPS2* enhanced MDC biogenesis (Fig. 1 D). To verify these preliminary results, we deleted *UPS1* and *UPS2*, and analyzed the lipid profiles and MDC biogenesis rates in these strains. Consistent with previous studies (Miyata et al., 2016; Connerth et al., 2012), deleting *UPS1* depleted cellular CL, while loss of *UPS2* resulted in reduced cellular PE abundance (Fig. S1 A). In line with the screen results, MDC biogenesis was impaired in *ups1*Δ cells in response to all known MDC inducers, including Rap, ConcA, and CHX (Fig. 2, B and C, Fig. S1 B). In contrast, even without the addition of MDC-inducing agents, 40% of *ups2*Δ mutant cells constitutively formed MDCs, and this percentage further increased when treated with MDC inducers (Fig. 2, B and C, Figure S1 B).

Previous studies showed that *UPS2* deletion rescues CL deficiency and mitochondrial import defects in *ups1*Δ mutants (Tamura et al., 2009), possibly through a Ups1-independent CL synthesis pathway activated by the ablation of mitochondrial PE production (Miyata et al., 2017). We constructed *ups1Δups2*Δ double mutants, and verified that their CL abundance was partially replenished compared with *ups1*Δ mutants (Fig. S1 A). Next, we performed MDC assays on *ups1Δups2*Δ double mutants. We found that this strain exhibited intermediate MDC formation compared to wild type, *ups1*Δ, and *ups2*Δ single mutants. Similar to *ups2*Δ, 30% of untreated *ups1Δups2*Δ cells formed MDCs (Fig. 2, B and C, Fig. S1 B). However, upon addition of MDC inducers, this percentage was not significantly increased and was lower than wild-type and *ups2*Δ cells—a phenotype that is similar to *ups1*Δ mutants (Fig. 2, B and C, Fig. S1 B). In addition, deletion of *MDM35*, a gene encoding a protein that binds to and stabilizes both Ups1 or Ups2 (Miyata et al., 2016; Potting et al., 2010), led to similar phospholipid composition (Fig. S1 A) and MDC biogenesis phenotypes (Fig. 2, B and C, Fig. S1 B) as observed in *ups1Δups2*Δ cells. These data support the results from the screen that *UPS1* and *UPS2* regulate MDC biogenesis in an inverse manner.

### Deficient mitochondrial CL synthesis inhibits MDC biogenesis

Reduced MDC formation in *ups1*Δ mutants suggests that CL may be required for MDC biogenesis. To test this, we analyzed MDC formation in additional mutants that perturb CL biosynthesis. In mitochondria, after PA is translocated from the OMM to the IMM by Ups1, it is converted to CL through several reactions catalyzed by enzymes localized in the IMM (Acoba et al., 2020; Tatsuta et al., 2014) (Fig. 3 A). We deleted non-essential genes encoding mitochondrial CL synthesis pathway components, including *GEP4*, which catalyzes the formation of phosphatidylglycerol (PG), an intermediate product of CL synthesis (Osman et al., 2010), as well as *CRD1*, the CL synthase that catalyzes CL formation from PG (Chang et al., 1998). Similar to *ups1*Δ cells, *gep4*Δ and *crd1*Δ strains were depleted of CL (Fig. S1 C) and showed impaired MDC biogenesis in response to Rap, ConcA, or CHX (Fig. 3, B and C, Fig. S1 D). In addition to pharmaceutical inducers, genetic activation of MDC formation through overexpressing mitochondrial SLC25A carrier proteins *AAC1* and *OAC1* (Schuler et al., 2021) was also prevented by impairing CL synthesis (Fig. S1 E). Our data indicate that CL is required for MDC formation.

**Figure 3.**
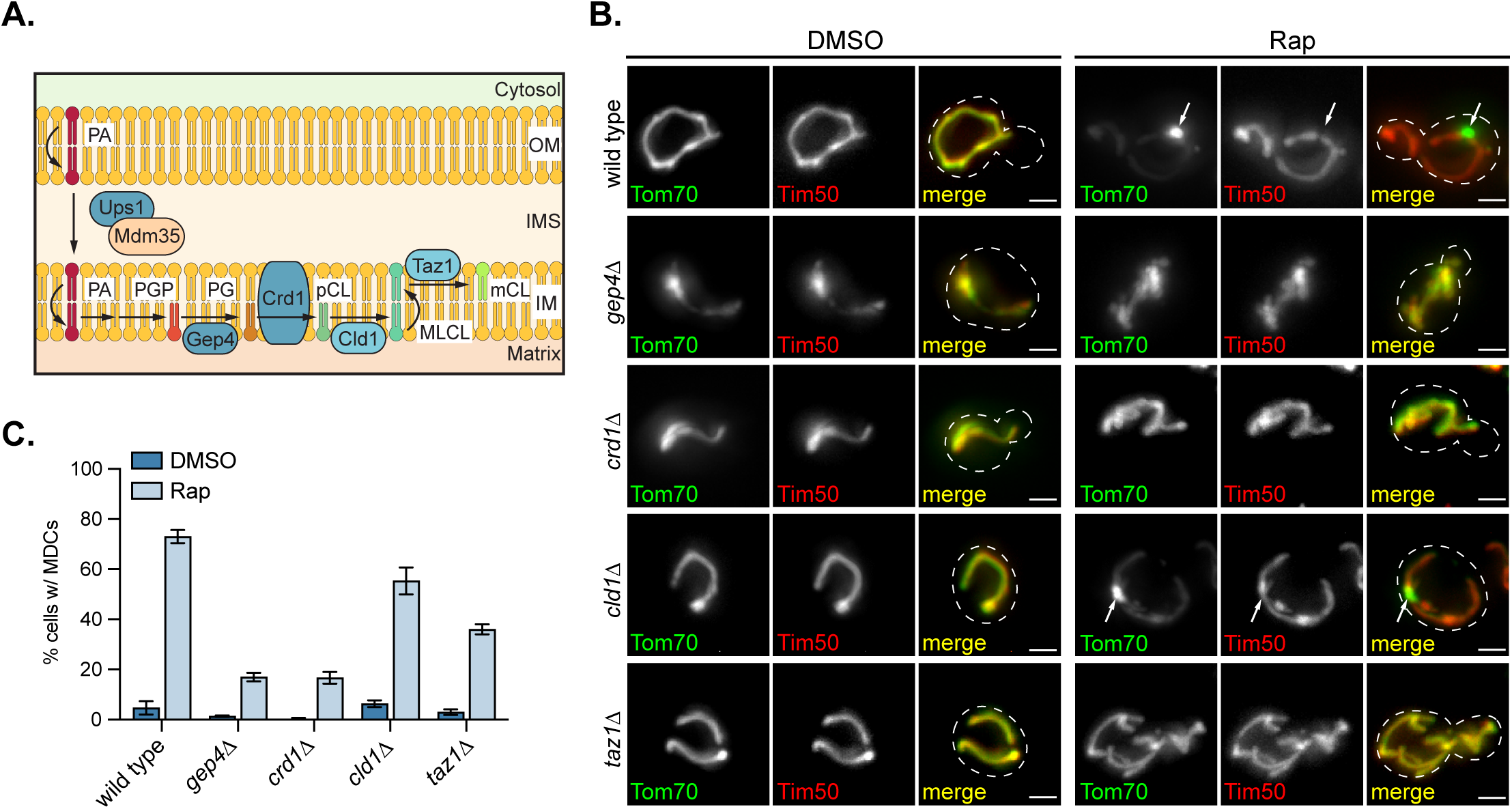
Deficiency in mitochondrial CL synthesis inhibits MDC biogenesis. **(A)** Model of the mitochondrial CL synthesis pathway. After the translocation of PA from the OMM to IMM by the Ups1-Mdm35 complex, CL is produced via several reaction steps. Gep4 catalyzes the synthesis of PG, an intermediate substrate for CL synthesis, and the CL synthase Crd1 converts PG to premature CL (pCL). Cld1 removes one acyl-chain from pCL to generate monolyso-CL (MLCL), followed by the re-addition of an acyl-chain to form mature CL (mCL) by Taz1. **(B)** Widefield images of wild-type cells or the indicated mutant yeast endogenously expressing Tom70-GFP and Tim50-mCh treated with DMSO or Rap for 2 h. White arrows mark positions of MDCs. Scale bar = 2 µm. **(C)** Quantification of (B) showing the percentage of cells with MDCs. N > 100 cells per replicate, error bars = SEM of three replicates.

After synthesis, CLs are further remodeled to generate mature CLs (Acoba et al., 2020). The maturation of CLs requires Cld1, which removes an acyl chain from premature CLs to form monolyso-CLs (Ye et al., 2014), and Taz1, which re-acylates monolyso-CLs to form mature CLs (Gu et al., 2004) (Fig. 3 A). To test whether CL remodeling affects MDC biogenesis, we assayed MDC formation in strains deleted for *CLD1* and *TAZ1*. We found *CLD1* deletion inhibited MDC formation induced by ConcA, and slightly impaired MDC biogenesis induced by Rap or CHX treatment (Fig. 3, B and C, Fig. S1 D). In comparison, deletion of *TAZ1* led to reduced MDC biogenesis in response to all MDC inducers (Fig. 3, B and C, Fig. S1 D). However, the inhibitory effect of *TAZ1* deletion was not as strong as deletion of *GEP4* or *CRD1*. These results suggest that CL remodeling has a mild impact on MDC biogenesis, but is not as critical as production of CL from PA.

### Defective mitochondrial PE synthesis constitutively activates MDC biogenesis

In addition to the CL synthesis pathway, we also investigated how deficient mitochondrial PE production on the IMM affects MDC biogenesis. Similar to *UPS2*, we identified *PSD1*, which encodes for the predominantly IMM and partially ER-localized decarboxylase that catalyzes PE synthesis from PS (Tatsuta et al., 2014; Friedman et al., 2018) (Fig. 4 A), was also identified as a potential negative regulator of MDC formation in the screen (Fig. 1 D). We then deleted *PSD1* and found that indeed, 35% of untreated *psd1*Δ cells constitutively formed MDCs (Fig. 4, B and C), indicating the MDC pathway is activated in the absence of mitochondrial PE synthesis.

**Figure 4.**
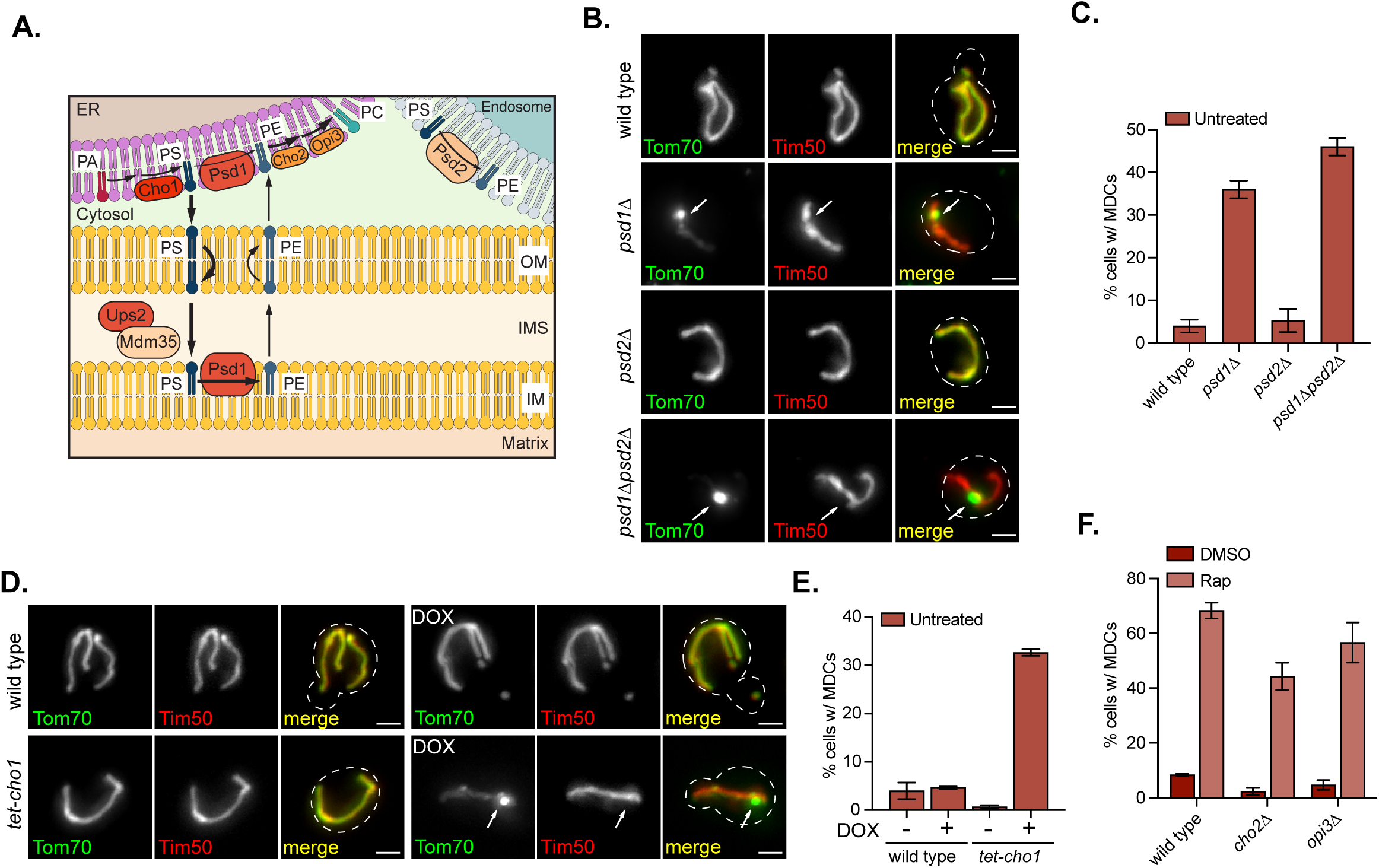
Defective mitochondrial PE synthesis constitutively activates MDC biogenesis. **(A)** Model of PE synthesis pathways by mitochondrial-localized Psd1 or endosomal-localized Psd2. PS, the substrates for PE synthesis, is synthesized from PA on the ER membrane. The last step of the reaction to generate PS is catalyzed by ER-localized Cho1. After transfer from the OMM to the IMM by the Ups2-Mdm35 complex, PS is converted to PE by an IMM decarboxylase Psd1. PE produced in mitochondria is further transferred to the ER for PC synthesis, which is mediated by Cho2 and Opi3 on the ER. An alternative PE synthesis pathway mediated by Psd2 is localized on the endosomal membrane. Psd1 can also dually localize to the ER membrane. **(B)** Widefield images of wild-type cells or the indicated mutant yeast endogenously expressing Tom70-GFP and Tim50-mCh. White arrows mark positions of MDCs. Scale bar = 2 µm. **(C)** Quantification of (B) showing the percentage of cells with MDCs. N > 100 cells per replicate, error bars = SEM of three replicates. **(D)** Widefield images of wild-type cells or *tet-cho1* mutants endogenously expressing Tom70-GFP and Tim50-mCh treated with DMSO or Rap for 2 h in the absence or presence of Doxycycline (Dox). White arrows mark positions of MDCs. Scale bar = 2 µm. **(E)** Quantification of (D) showing the percentage of cells with MDCs. N > 100 cells per replicate, error bars = SEM of three replicates. **(F)** Quantification of MDC formation in wild-type cells and the indicated mutant yeast treated with DMSO or Rap for 2 h. N > 100 cells per replicate, error bars = SEM of three replicates.

In yeast, Psd1-dependent PE synthesis is not the only pathway for PE production. In *psd1*Δ cells, PE produced by other cellular pathways partially supplies mitochondrial PE (Trotter and Voelker, 1995; Birner et al., 2001; Bürgermeister et al., 2004; Gulshan et al., 2010; Gibellini and Smith, 2010; Acoba et al., 2020). From previous studies (Schuler et al., 2016) and as verified here, *psd1*Δ mutants only contain slightly reduced whole-cell PE levels compared with wild-type cells (Fig. S2 A). To test whether MDC biogenesis is exacerbated by depletion of redundant cellular PE production, we deleted *PSD2*, which catalyzes PE synthesis on endosomal membranes (Tatsuta et al., 2014) (Fig. 4 A). The deletion of *PSD2* itself did not affect total cellular PE levels (Fig. S2 A) or MDC biogenesis (Fig. 4, B and C). However, whole-cell PE abundance was severely reduced in *psd1Δpsd2*Δ double mutants (Fig. S2 A). We found that 45% of *psd1Δpsd2*Δ cells constitutively formed MDCs, which is slightly higher than the rate in *psd1*Δ single mutants (Fig. 4, B and C). We also tested whether adding MDC-inducing agents further increased MDC biogenesis in *psd1*Δ and *psd1Δpsd2*Δ mutants. Although the percentages of cells with MDCs was increased compared with untreated *psd1*Δ and *psd1Δpsd2*Δ mutants, these mutants failed to outcompete wild-type cells in MDC formation triggered by Rap, ConcA, or CHX (Fig. S2 B). In contrast, MDC biogenesis induced by overexpression of *AAC1* and *OAC1* (Schuler et al., 2021) was further enhanced by *PSD1* deletion (Fig. S2 C).

Constitutive MDC biogenesis in cells with deficient mitochondrial PE synthesis could be a result of the accumulation of the PE substrate PS on the OMM or impaired phosphatidylcholine (PC) synthesis from PE (Miyata et al., 2017; Tatsuta et al., 2014) (Fig. 4 A). To test these hypotheses, we first knocked down *CHO1*, the enzyme that catalyzes the last step of PS synthesis on the ER membrane (Miyata et al., 2017). In response to doxycycline (Dox)-dependent *Tet*-induced inhibition, *tet-CHO1* mutants showed depleted *CHO1* mRNA abundance (Fig. S2 D). Interestingly, inhibiting *CHO1* expression caused constitutive MDC biogenesis similar to *ups2*Δ and *psd1*Δ strains (Fig. 4 D, Fig. S2 E), indicating that MDC biogenesis in *ups2*Δ and *psd1*Δ mutants is not a consequence of PS accumulation on the OMM. Second, to test whether inhibiting mitochondrial PE production induces MDC formation by limiting substrates for PC synthesis on the ER membrane (Tatsuta et al., 2014), we deleted *CHO2* and *OPI3*, two enzymes that catalyze PC generation from PE (Kodaki and Yamashita, 1987, 1989). Specifically, Cho2 drives the first step of PE methylation to generate monomethyl-PE (MMPE), and Opi3 promotes all three methylation steps to synthesize MMPE, dimyristoyl-PE (DMPE), and PC sequentially (Kodaki and Yamashita, 1987, 1989). Unlike *ups2*Δ and *psd1*Δ mutants, untreated *cho2*Δ and *opi3*Δ mutants did not form MDCs constitutively (Fig. 4 F). On the contrary, deletions of *CHO2* and *OPI3* lead to slightly decreased MDC biogenesis in response to Rap or ConcA compared to wild-type cells (Fig. 4 F, Fig. S2 F), indicating that the MDC biogenesis triggered by deletion of *UPS2* and *PSD1* is not caused by impaired PC production. Interestingly, in *cho2*Δ and *opi3*Δ mutants that fail to synthesize PC from PE, mitochondrial PE is elevated (Schuler et al., 2016), which likely explains the modest reduction in MDC formation. Taken together, our results suggest that lack of mitochondrial PE, but not alterations in the abundance of PS or PC, stimulates MDC biogenesis.

### *PSD1* deletion partially restores MDC biogenesis in ERMES/Gem1 mutants

We previously showed that proteins at ER-mitochondria contacts are required for MDC biogenesis (English et al., 2020). Deletion of individual ERMES components or its regulatory factor *GEM1* prevents MDC biogenesis (English et al., 2020). However, at that time, it was unclear whether mitochondrial phospholipid homeostasis, which is one of the major functions of the ERMES complex (Kornmann et al., 2009; Tatsuta et al., 2014; AhYoung et al., 2015; Kojima et al., 2016; Jeong et al., 2017), regulates MDC biogenesis. Given that defective mitochondrial PE synthesis constitutively activates MDC biogenesis, we tested whether PE depletion could bypass the requirement for ERMES/Gem1 in MDC formation. To do this, we deleted *UPS2* or *PSD1* in ERMES mutants expressing *VPS13(D716H)*, which is a mutant version of *VPS13* that has been shown to rescue mitochondrial morphology, but not ERMES complex assembly (Lang et al., 2015) or MDC biogenesis (English et al., 2020) in strains lacking individual ERMES components. In the absence of MDC-inducing agents, *PSD1* deletion led to constitutive MDC formation in 25% of *mmm1Δ VPS13(D716H)* cells, which lack the ERMES component Mmm1. This percentage increased to 60% with the addition of Rap or CHX, but not ConcA (Fig. 5, A and B, Fig. S2 G). Deleting *UPS2* in *mmm1Δ VPS13(D716H)* cells only moderately restored MDC biogenesis when cells were treated with Rap or CHX (Fig. 5, A and B, Fig. S2 G). We also tested whether deletion of *UPS2* or *PSD1* restored MDC formation in mutants lacking *GEM1*, the GTPase that regulates the ERMES complex (Kornmann et al., 2011). Deletion of *GEM1* is one of the most robust MDC inhibitors identified so far as all common suppressors of ERMES mutants fail to rescue MDC formation in *gem1*Δ mutants (English et al., 2020). We found that with the addition of Rap, ConcA, or CHX, *PSD1* deletion rescued MDC formation in less than 20% of *gem1*Δ mutants (Fig. 5, C and D, Fig. S2 H), and *UPS2* deletion was incapable of rescuing MDC formation in *gem1*Δ mutants (Fig. 5, C and D, Fig. S2 H), indicating that Ups2 and Psd1 somewhat differ in their abilities to regulate MDC biogenesis in the absence of the ER-mitochondria contacts. These results suggest that genetically inhibiting mitochondrial PE synthesis can bypass the requirement for ERMES in the MDC pathway, raising the idea that the role of the ERMES complex in MDC formation is likely linked to its function in phospholipid homeostasis. Because restoration of MDC formation in *gem1*Δ mutants was not as robust, it is possible that Gem1 plays additional roles in MDC biogenesis.

**Figure 5.**
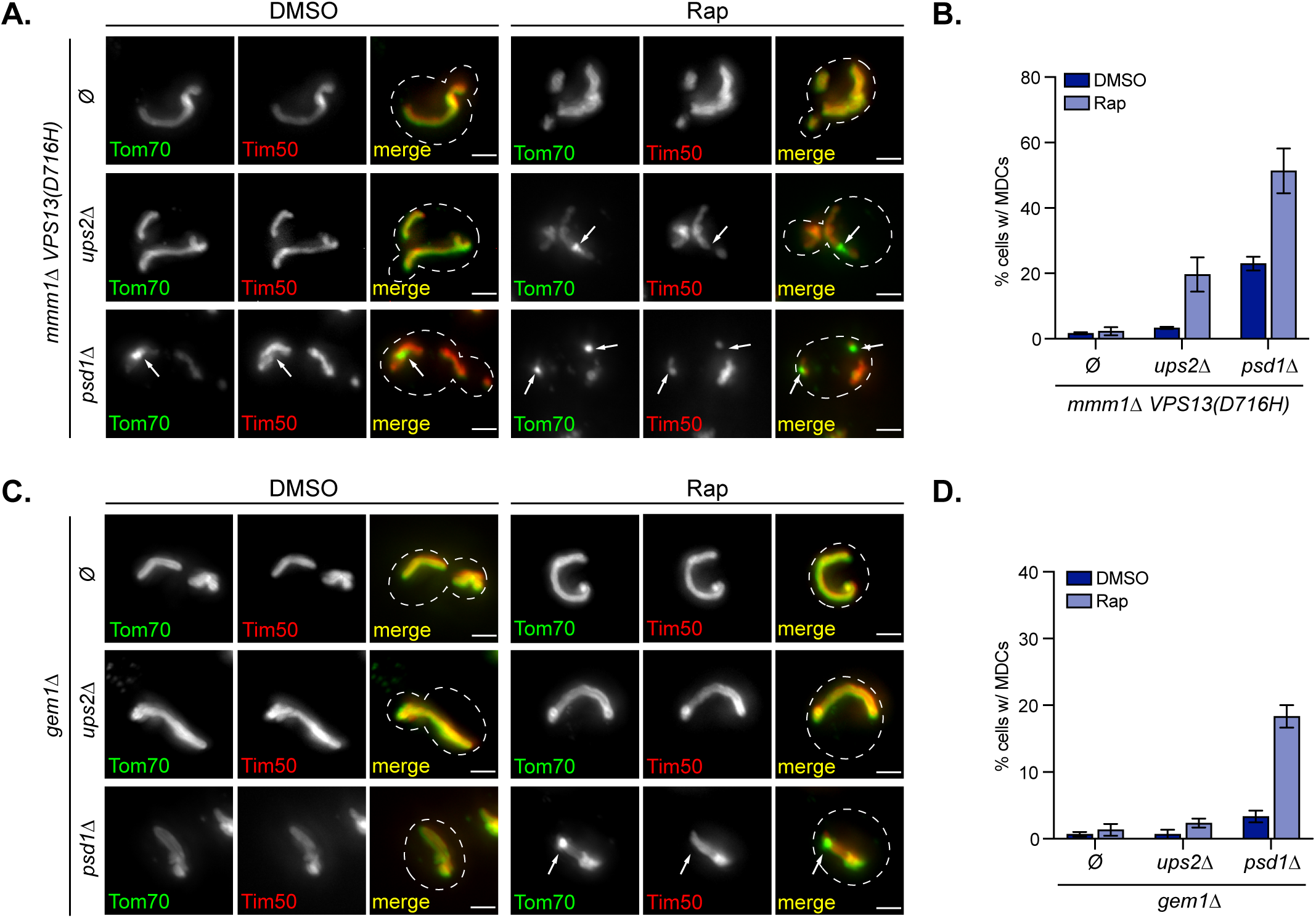
*PSD1* deletion partially restores MDC biogenesis in ERMES/Gem1 mutants. **(A)** Widefield images of *mmm1Δ VPS13(D716H)* cells or *mmm1Δ VPS13(D716H)* cells with the indicated gene deletions endogenously expressing Tom70-GFP and Tim50-mCh treated with DMSO or Rap for 2 h. Ø, no gene is deleted. White arrows mark positions of MDCs. Scale bar = 2 µm. **(B)** Quantification of (A) showing the percentage of cells with MDCs. N > 100 cells per replicate, error bars = SEM of three replicates. **(C)** Widefield images of *gem1*Δ cells or *gem1*Δ cells with the indicated gene deletions endogenously expressing Tom70-GFP and Tim50-mCh treated with DMSO or Rap for 2 h. Ø, no gene is deleted. White arrows mark positions of MDCs. Scale bar = 2 µm. **(D)** Quantification of (C) showing the percentage of cells with MDCs. N > 100 cells per replicate, error bars = SEM of three replicates.

### Cellular and mitochondrial PE declines in response to various MDC-inducing agents

In addition to investigating how genetically altering mitochondrial phospholipid synthesis affects MDC biogenesis, we also analyzed whether cellular and mitochondrial lipid composition changes in response to known MDC-inducing agents, including ConcA, Rap, and CHX. Through mass spectrometry-based lipidomic analysis on whole-cell lipid extracts, we found that all three MDC inducers triggered a decrease in PS and PE abundance (Fig. 6 A). PC levels were unaffected by ConcA and slightly reduced by Rap and CHX treatment (Fig. 6 A). ConcA, but not the other two inducers, dramatically increased cellular PG abundance (Fig. 6 A). Whole-cell CL levels, however, were not significantly affected in response to all three inducers (Fig. 6 A).

**Figure 6.**
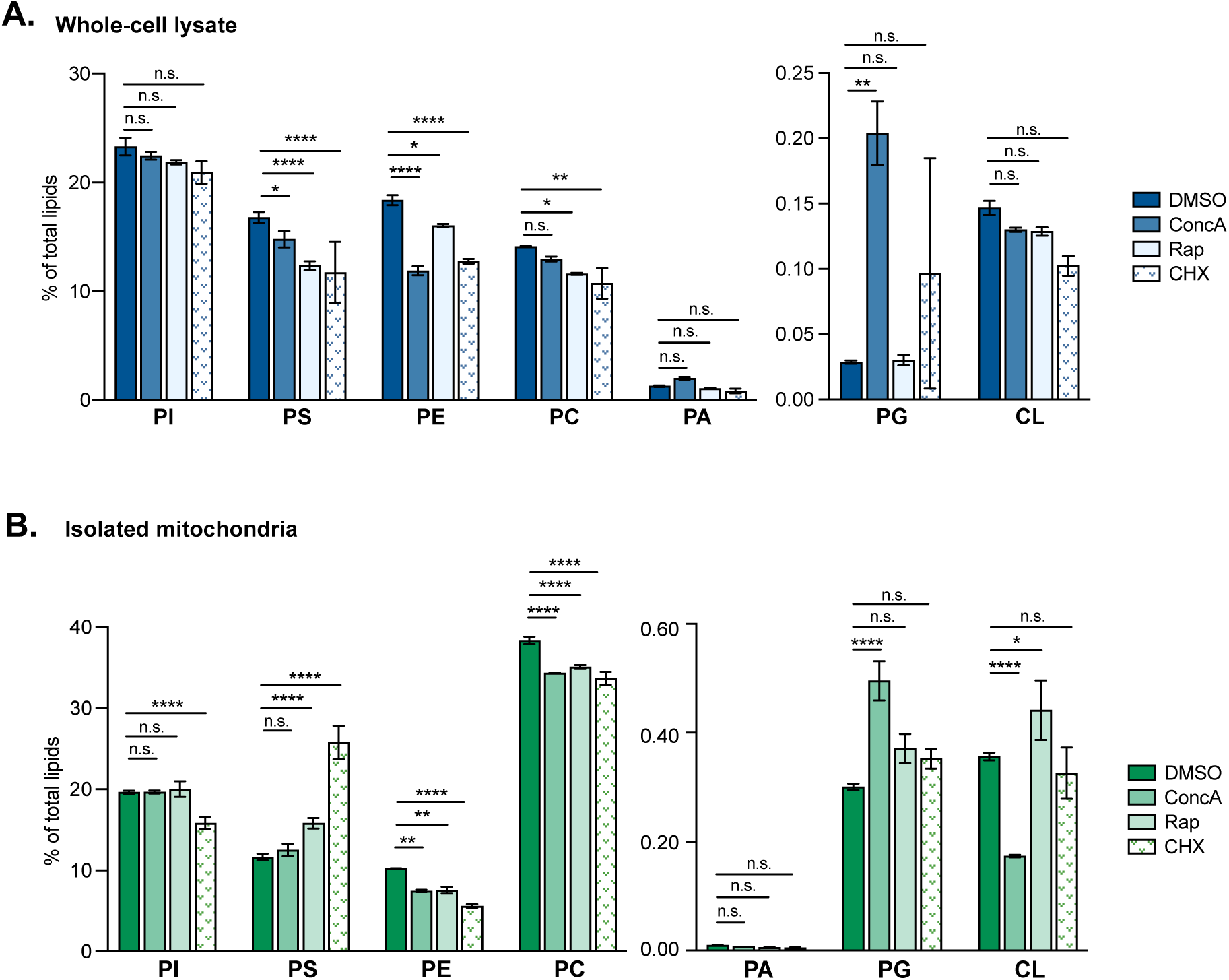
Cellular and mitochondrial PE levels decline under MDC-inducing conditions. **(A and B)** The relative amounts of the indicated phospholipids in whole-cell lysates (A) or crude mitochondria preparations (B) isolated from yeast cells treated with DMSO, ConcA, Rap, or CHX for 2 h detected by mass spectrometry-based lipidomic analysis. Amounts of each lipid relative to total lipids were determined. Error bars = SEM of three replicates. Statistical comparison shows difference to the corresponding DMSO control. n.s., not significant, *p<0.0332, **p<0.0021, ***p<0.0002, ****p<0.0001, 2-way ANOVA with Holm-ŠÍdák test.

We also observed lower PE in mitochondria isolated from the cells treated with ConcA, Rap, or CHX (Fig. 6 B). Mitochondrial PS levels, in contrast to whole-cell PS, were unaffected or increased (Fig. 6 B). With all three inducers, mitochondrial PC abundance decreased (Fig. 6 B), which could be explained by the lack of PE, the substrate for PC synthesis (Tatsuta et al., 2014). Consistent with the observed elevation in total cellular PG levels, mitochondrial PG abundance was also significantly increased by ConcA, but not Rap or CHX (Fig. 6 B). The mitochondrial CL levels exhibit distinct alterations in response to different MDC-inducing agents, which were decreased by ConcA, increased by Rap, and unaffected by CHX (Fig. 6 B).

Our genetic results showed that depleting CL inhibits MDC biogenesis (Fig. 3). However, we did not observe any consistent changes in CL abundance across different MDC inducers, suggesting that the generation of MDCs requires the presence of CL, but is unlikely initiated by an acute alteration in CL levels. In contrast, for all three MDC-inducing agents, we observed a decline in both cellular and mitochondrial PE abundance, which is consistent with the result that MDC biogenesis is activated by genetically impairing mitochondrial PE synthesis (Fig. 2, B and C, Fig. 4, B and C). These results raise the possibility that reduced mitochondrial PE levels may trigger MDC formation.

To further explore the connection between PE abundance and the MDC pathway, we next analyzed whether known mutants with up- or down-regulated MDC biogenesis exhibit differences in whole-cell PE levels upon Rap treatment. We found that in mutants with deficient CL synthesis, including *ups1*Δ, *gep4*Δ and *crd1*Δ strains, their PE depletion by Rap was similar to wild-type cells (Fig. S3, A and B). Thus, it is unlikely that the requirement for CL in MDC biogenesis is linked to changes in PE. In contrast, mutant strains with defective mitochondrial PE production, including *ups2*Δ, *ups1Δups2*Δ, *mdm35*Δ, and *psd1*Δ, which have constitutively elevated MDC formation (Fig. 2, B and C, Fig. 3, B and C), exhibited an attenuated decrease in cellular PE levels induced by Rap compared with wild-type cells (Fig. S3, A and C). This suggests that genetically inhibiting mitochondrial PE synthesis and Rap treatment share some redundancy in activating MDC formation. Interestingly, *psd2*Δ mutants showed dramatic PE depletion compared with wild-type cells, and this depletion was not further exacerbated in *psd1Δpsd2*Δ mutants (Fig. S3 C), indicating a decline of PE synthesis by Psd1, but not Psd2, is triggered by MDC-inducing stress.

In addition to mitochondrial phospholipid synthesis pathway mutants, we tested whether lack of *GEM1*, a regulatory protein of ER-mitochondria contacts (Kornmann et al., 2011) that is required for MDC formation (English et al., 2020), affects PE abundance changes in response to Rap induction. Indeed, we found PE decline was blunted in *gem1*Δ cells (Fig. S3 D), which coincides with the result that deletion of *GEM1* prevents MDC biogenesis. Taken together, our data suggests a significant regulatory role of PE abundance in MDC biogenesis.

### Overexpressing *UPS2* and *PSD1* inhibits MDC biogenesis

To further investigate whether PE decline is required for MDC biogenesis more directly, we tested whether genetically upregulating mitochondrial PE synthesis pathway components could block MDC formation induced by Rap, CHX, and ConcA. Indeed, overexpressing *UPS2* or *PSD1* inhibited MDC formation in response to ConcA (Fig. 7, A and B). In cells treated with Rap or CHX, MDC biogenesis was prevented by *UPS2* overexpression, and was slightly impaired by *PSD1* overexpression (Fig. S3 E). Analysis of the lipid composition of cells overexpressing *UPS2* revealed that PE decline was blunted in these cells (Fig. 7 C). Cells overexpressing *PSD1*, on the other hand, exhibited higher baseline PE levels than wild-type cells and showed a moderate increase in CL (Fig. 7 C). However, PE levels still declined upon ConcA treatment in these cells, albeit to a slightly higher final level than in wild-type cells (Fig. 7 C). Notably, levels of MMPE and DMPE were elevated in both untreated and treated *PSD1* overexpressing cells (Fig. S3 F), suggesting these downstream PE products may also play a role in MDC regulation. Altogether, these results indicate that boosting the expression of mitochondrial PE synthesis pathway components prevents MDC formation.

**Figure 7.**
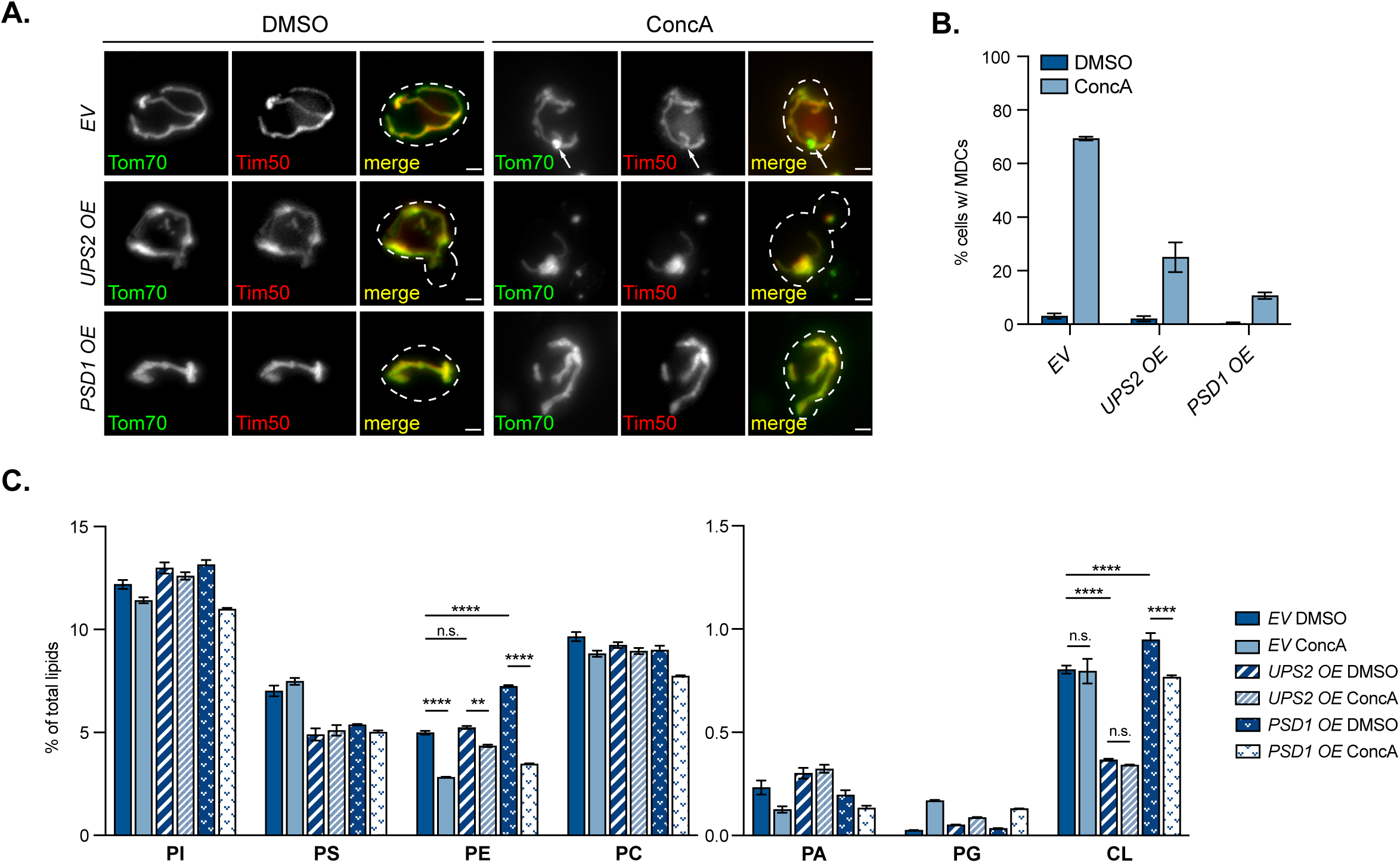
Overexpressing *UPS2* or *PSD1* suppresses MDC formation. **(A)** Widefield images of yeast cells with genomic integration of an empty vector (*EV*), *UPS2* overexpressing vector (*UPS2 OE*), or *PSD1* overexpressing vector (*PSD1 OE*) endogenously expressing Tom70-GFP and Tim50-mCh treated with DMSO or ConcA for 2 h. White arrows marks positions of MDCs. Scale bar = 2 µm. **(B)** Quantification of (A) showing the percentage of cells with the indicated integrated vectors with MDCs. N > 100 cells per replicate, error bars = SEM of three replicates. **(C)** The relative amounts of the indicated phospholipids in whole-cell lysates from yeast cells with genomic integration of an empty vector (*EV*), *UPS2* overexpressing vector (*UPS2 OE*), or *PSD1* overexpressing vector (*PSD1 OE*) treated with DMSO or ConcA for 2 h detected by mass spectrometry-based lipidomic analysis. Amounts of each lipid relative to total lipids were determined. Error bars = SEM of three replicates. Statistical comparison shows difference to the corresponding DMSO control. n.s., not significant, *p<0.0332, **p<0.0021, ***p<0.0002, ****p<0.0001, 2-way ANOVA with Holm-ŠÍdák test.

### Depletion of PE abundance correlates with amino acid levels

Next, we further explored the hypothesis that PE depletion triggers MDC formation downstream of amino acid elevation caused by Rap, CHX, and ConcA. We previously reported that reducing or eliminating amino acids from the media in which yeast cells are cultured completely blocks ConcA-induced MDCs, and impairs Rap- and CHX-induced MDCs (Schuler et al., 2021) (Fig. 8 A, Fig. S4 A). Thus, we tested whether the depletion of PE in response to ConcA was also dependent on amino acid levels in the media. Indeed, although we still observed a decrease in cellular PE abundance with ConcA treatment when cells are grown in synthetic media containing low levels of amino acids or in minimal media that completely exclude amino acids, the extent of their PE depletion was attenuated compared with cells grown in amino acid-enriched media (Fig. 8 B, Figure S4 B-D).

**Figure 8.**
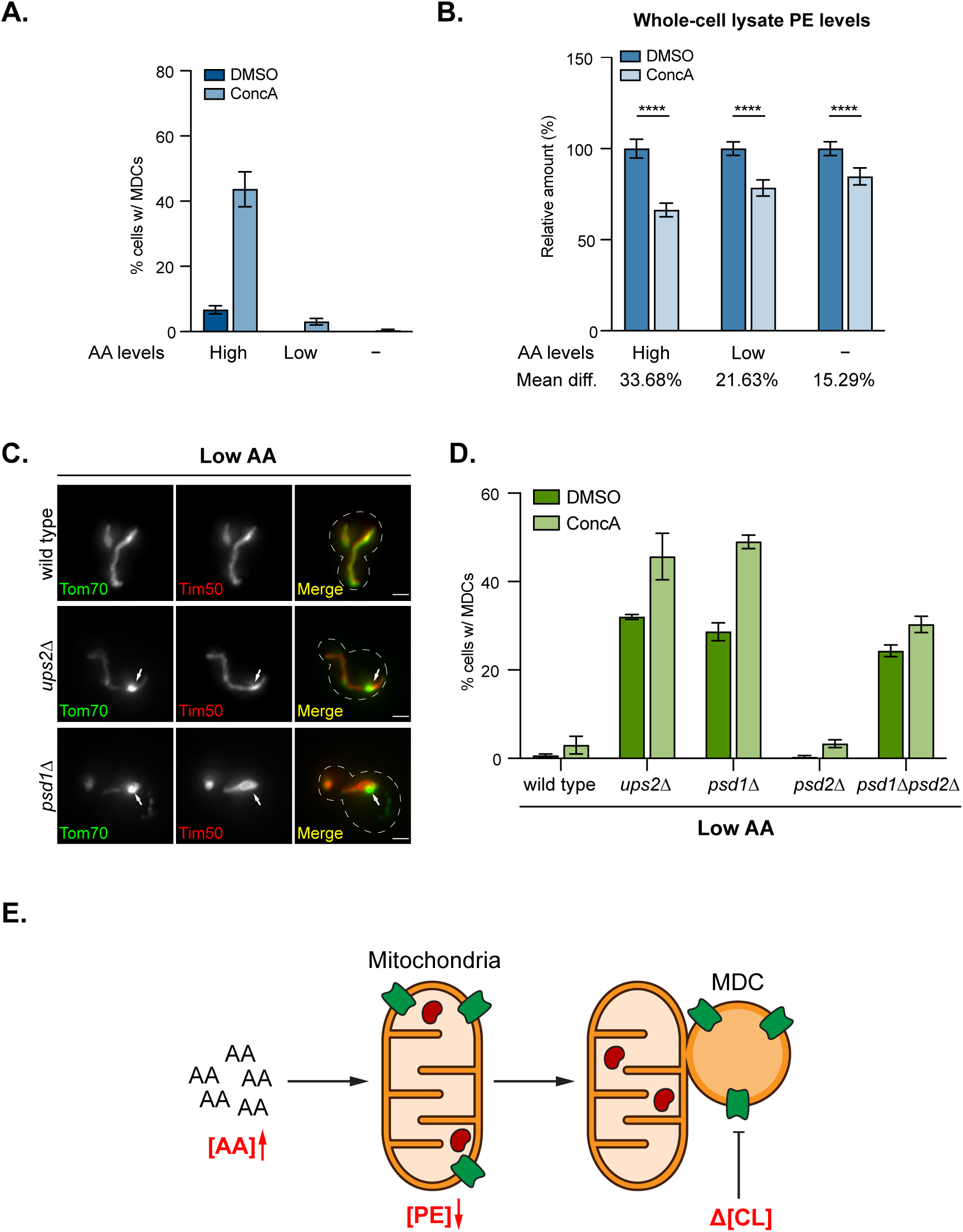
PE decline activates MDC formation downstream of amino acid stress. **(A)** Quantification of MDC formation in yeast cells grown in amino acid-rich media, synthetic media that contains low levels of amino acid, and minimal media that exclude amino acids (indicated as High, Low, and −, respectively) treated with DMSO or ConcA for 2 h. N > 100 cells per replicate, error bars = SEM of three replicates. **(B)** The relative amounts of PE in whole-cell lysates of yeast cells grown in amino acid-rich media, synthetic media that contains low levels of amino acid, and minimal media that exclude amino acids (indicated as High, Low, and −, respectively) treated with DMSO or ConcA for 2 h. The amounts of PE in DMSO-treated cells were set to 100%. Mean diff. = mean difference between the amounts of PE in DMSO- and ConcA-treated cells cultured in the indicated media. Error bars = SEM of three replicates. Statistical comparison shows difference to the corresponding DMSO control. n.s., not significant, *p<0.0332, **p<0.0021, ***p<0.0002, ****p<0.0001, 2-way ANOVA with Holm-ŠÍdák test. **(C)** Widefield images of wild-type cells or the indicated mutant yeast endogenously expressing Tom70-GFP and Tim50-mCh grown in media containing low levels of amino acids treated with DMSO or ConcA for 2 h. White arrows marks positions of MDCs. Scale bar = 2 µm. **(D)** Quantification of MDC formation in wild-type cells or the indicated mutant cells treated with DMSO or ConcA. N > 100 cells per replicate, error bars = SEM of three replicates. **(E)** Model of MDC formation regulated by CL and PE in response to MDC-inducing amino acid stress. Amino acid elevation triggers PE decline, which activates MDC formation. Depletion of CL production prevents MDC formation caused by amino acid stress.

### Deletion of *UPS2* or *PSD1* restores MDC formation in low amino acid media

Given that reducing amino acid abundance in the media attenuates PE depletion in response to MDC-inducing agents, we next tested whether genetically depleting mitochondrial PE could re-activate MDC biogenesis in cells cultured in media containing low levels of amino acids. We found that 30% of *ups2*Δ and *psd1*Δ mutants constitutively formed MDCs in synthetic low amino acid media (Fig. 8, C and D). Interestingly, this percentage is comparable to *ups2*Δ and *psd1*Δ mutants cultured in amino acid-rich media (Fig. 2 C, Fig. 4 C), suggesting amino acid abundance is dispensable for activating MDC biogenesis induced by defective mitochondrial PE production. The percentages of *ups2*Δ and *psd1*Δ mutants that formed MDCs was further increased by the addition of MDC-inducing agents (Fig. 8 D, Fig. S4 E). In contrast to high amino acid media, deleting *PSD2* in *psd1*Δ mutants did not exacerbate MDC formation in the absence or presence of MDC inducers (Fig. 8 D, Fig. S4 E), indicating that for *psd1*Δ mutants grown in low amino acid media, mitochondria are less likely to obtain PE synthesized by Psd2. These results suggest that PE depletion induces MDC biogenesis independently of amino acid levels. In combination with our other observations, we propose that PE decline may act as a signal for MDC formation downstream of amino acid overabundance.

## DISCUSSION

We previously showed that the MDC pathway is activated by elevated intracellular amino acid levels (Schuler et al., 2021), and that MDCs form at ER-mitochondria contacts in an ERMES/Gem1-dependent manner (English et al., 2020). However, it remained unclear from these studies how amino acid changes initiate MDC biogenesis, as well as whether phospholipid transport, which is a major function of the ERMES complex (Tatsuta et al., 2014), affects the generation of MDCs. In this study, we used an unbiased genome-wide imaging screen to identify genetic regulators of the MDC pathway. This approach identified a number of gene deletions that impact MDC biogenesis, suggesting MDC formation rates are influenced by a variety of cellular processes. Amongst gene hits, we focused on further characterization of a set of factors linked to mitochondrial phospholipid homeostasis, given the known role for ERMES and ER-mitochondria contact sites in MDC formation (English et al., 2020). Through a combination of genetics and mass-spectrometry-based lipidomic analyses, we uncovered distinct roles for CL and PE in MDC formation (Fig. 8 E). We found that CL is required for MDC biogenesis, as deletion of lipid transporters and enzymes in the CL synthesis pathway, including *UPS1*, *GEP4*, and *CRD1*, prevented MDC formation (Fig. 2, B and C, Fig. 3, B and C). In contrast, we showed that PE levels declined in an amino acid-dependent manner upon treatment of cells with MDC inducers Rap, CHX, and ConcA (Fig. 6, A and B, Fig. 8, A and B), found that genetic impairment of mitochondrial PE synthesis by deleting *UPS2* or *PSD1* was sufficient to trigger MDC formation regardless of the amino acid abundance in the media (Fig. 2, B and C, Fig. 4, B-E, Fig. 8, C and D), and showed that overexpression of PE synthesis components *UPS2* and *PSD1* blunted MDC formation (Fig. 7, A and B). Based on these results, we propose that PE decline triggers MDC formation downstream of amino acid overabundance stress (Fig. 8 E).

Currently, it remains unclear how CL and PE regulate MDC formation, and why the two lipids, which are often thought to have redundant and overlapping roles in supporting various aspects of mitochondrial biology (Acoba et al., 2020; Basu Ball et al., 2018), would have opposing impacts on MDC biogenesis. Both CL and PE are critical for mitochondrial respiratory chain activity, mitochondrial membrane potential, and mitochondrial import (Acoba et al., 2020; Gohil and Greenberg, 2009). Deleting both *CRD1* and *PSD1*, which encode IMM enzymes that catalyze CL and PE synthesis respectively, leads to synthetic lethality in yeast (Gohil et al., 2005). In contrast, PE and CL appear to have distinct roles in MDC biogenesis. Further studies are necessary to understand why cells that lack CL do not form MDCs. It is possible CL plays a direct role in forming the MDC membrane structure. It is also equally likely that CL regulates MDC formation by perturbing an aspect of mitochondrial function critical for MDC biogenesis, or by somehow bypassing the need to form MDCs.

Likewise, due to the many impacts of PE on mitochondrial homeostasis, we have not yet identified a precise role for PE in MDC regulation. Nevertheless, our results provide a step forward in understanding how amino acid surplus could lead to alterations in mitochondrial structure and composition via the MDC pathway. Upon treatment with three different inhibitors that trigger intracellular amino acid accumulation, PE levels decline in the mitochondria. Our data show that removal of amino acids prevents PE decline (Fig. 8, A and B), suggesting that elevated amino acids, but not vacuole impairment or translation inhibition per se, lead to decreased PE. Moreover, genetic depletion of PE is epistatic to amino acids, as deleting PE synthesis machinery triggers MDCs regardless of amino acid status in the media (Fig. 8, C and D). These results suggest that PE depletion likely activates MDCs downstream of amino acid stress.

Interestingly, the concept of PE functioning as a lipid-based signal downstream of amino acid perturbation to drive MDC biogenesis exhibits strong similarities to a recently published study in mammals, which found that alterations in mitochondrial PE levels stimulate mitochondrial remodeling during hypoxia or impairment of the mTOR pathway (MacVicar et al., 2019). In that study, the authors found that mTOR inhibition decreased PE levels by stimulating the activity of LIPIN1, which drives the conversion of PA to diacylglycerol and triacylglycerol, thus lowering phospholipid levels. They also showed that a decline in PE triggered changes in mitochondrial protein abundance by activating the mitochondrial-localized protease, YME1L1 (MacVicar et al., 2019). In our current study, we also found that impairment of mTOR signaling by treatment of cells with Rap lowers PE and stimulates MDC formation. However, we also observed that CHX treatment, which activates mTOR signaling (Beugnet et al., 2003), also leads to lower PE levels, as does treatment with ConcA. Thus, while interesting parallels exist between the two organisms regarding the concept of PE as a signaling lipid downstream of cellular stress, it appears that the signaling relays may differ somewhat in yeast and mammals. Consistent with this idea, we found that deletion of the LIPIN1 homolog, *PAH1* (Fakas et al., 2011; Han et al., 2006), affects MDC biogenesis distinctly in response to different MDC-inducing agents. The *pah1*Δ strains showed enhanced MDC formation in vehicle- and ConcA-treated cells, unaltered MDC formation induced by Rap, and moderately inhibited CHX-triggered MDC biogenesis (Fig. S4 F). Thus, it remains to be determined how Rap, ConcA, and CHX promote PE decline in yeast. It is also unclear how PE decline stimulates MDC biogenesis. Of note, our preliminary screen results suggest that several mitochondrial proteases, including the YME1L1 homolog Yme1(Leonhard et al., 1999; Potting et al., 2010; Song et al., 2007; MacVicar et al., 2019), impact MDC formation (Fig. 1 D). Given the impact of PE levels on Yme1L1 activity in mammals (MacVicar et al., 2019), it will be interesting to determine in future studies whether a PE-protease axis controls MDC biogenesis in yeast. Overall, the results of our studies presented here enhance our understanding of the MDC pathway and its regulation, and further support the concept of PE functioning as key signaling node coupling cellular metabolic status to mitochondrial composition and function.

## MATERIALS AND METHODS

### Reagents

Plasmids and chemicals used in this study are listed in Table S2.

### Yeast strains

All yeast strains are derivatives of *Saccharomyces cerevisiae* S288c (BY) (Baker Brachmann et al., 1998) and are listed in Table S3. Strains expressing fluorescently tagged *TOM70* and *TIM50* from their native loci were created by one step PCR-mediated C-terminal endogenous epitope tagging using standard techniques and oligo pairs listed in Table S4. Plasmid templates for fluorescent epitope tagging were from the pKT series of vectors (Sheff and Thorn, 2004). Correct integrations were confirmed by correctly localized expression of the fluorophore by microscopy. For strains bearing deletion of *UPS1, UPS2, MDM35, GEP4, CRD1, CLD1, TAZ1, PSD1, PSD2, CHO2, OPI3,* or *GEM1,* one copy of the gene was deleted by one step PCR-mediated gene replacement with the indicated selection cassette using standard techniques and oligo pairs listed in Table S4 to create a heterozygous diploid, which was subsequently sporulated to obtain haploid mutants. Plasmid templates for gene replacement were from the pRS series of vectors (Sikorski and Hieter, 1989). Correct insertion of the selection cassette into the target gene was confirmed by colony PCR across the chromosomal insertion site. Yeast strains constitutively expressing *AAC1, OAC1, UPS2, or PSD1* from the GPD promoter were generated by integration of the expression cassette into yeast chr I (199456-199457) as previously described (Hughes and Gottschling, 2012). Plasmids for integration of the GPD-driven expression cassette are described in the Table S2. Correct insertion of the expression cassette into chr I was confirmed by colony PCR across the chromosomal insertion site.

Wild-type yeast strains AHY12872 and AHY11976, which were rendered prototrophic with pHLUM (Mülleder et al., 2012) (see Table S3) to prevent complications caused by amino acid auxotrophies in the BY strain background, were used to quantify amino acid dependencies of MDC formation and for analysis of whole cell lipid levels. Wild-type yeast strain AHY2318 was used for mating with the yeast deletion collection to create a new deletion library containing fluorescently tagged *TOM70* and *TIM50* for the MDC screen. AHY6760, AHY7019, AHY7021, AHY9543, AHY10568, AHY11351, AHY11353, AHY11357, AHY11359, AHY11367, AHY11435, AHY11438, AHY11442, AHY11533, AHY11634, AHY11766, AHY11768, AHY11966, AHY11990, AHY12011, AHY12518, AHY12775, AHY12803, AHY13656, AHY14155, AHY14157, AHY14226, AHY14228, AHY14230, AHY14232, AHY14234, and AHY14261 were used for quantification of MDC formation using widefield microscopy. Lipid analysis was performed in BY4741, AHY4230, AHY11362, AHY11370, AHY11440, AHY11533, AHY11636, AHY11874, AHY11964, AHY12516, AHY12736, and AHY12801. A complete list of all strains used in this manuscript can be found in Table S3.

### Yeast cell culture and media

Yeast cells were grown exponentially for 15 h at 30°C to a density of 4-8 ×10^6^ cells/ml before the start of all experiments described in the paper, including MDC assays and sample preparation for lipidomic analysis. This period of overnight log-phase growth was carried out to ensure mitochondrial uniformity across the cell population and is essential for consistent MDC activation. Unless otherwise indicated, cells were cultured in YPAD medium (1% yeast extract, 2% peptone, 0.005% adenine, and 2% glucose), which is the high amino acid media used for amino acid dependency experiments. For amino acid dependency experiments, SD medium (low amino acid medium) (0.67% yeast nitrogen base without amino acids, 2% glucose, supplemented nutrients 0.074 g/liter each adenine, alanine, arginine, asparagine, aspartic acid, cysteine, glutamic acid, glutamine, glycine, histidine, myo-inositol, isoleucine, lysine, methionine, phenylalanine, proline, serine, threonine, tryptophan, tyrosine, uracil, valine, 0.369 g/liter leucine, 0.007 g/liter para-aminobenzoic acid), or minimal medium (no amino acid medium) (0.67% yeast nitrogen base without amino acids, 2% glucose) were used. Drugs were added to cultures at final concentrations of 500 nM for Concanamycin A, 10 μg/ml for Cycloheximide, 20 mg/ml for Doxycycline, or 200 nM for Rapamycin.

### Plasmids

Plasmids used in this study are listed in the Table S2. pHLUM, a yeast plasmid expressing multiple auxotrophic marker genes from their endogenous promoters, was obtained from Addgene (#40276) (Mülleder et al., 2012). Plasmids for GPD-driven expression of *AAC1, OAC1, UPS2,* or *PSD1* were generated by gateway mediated transfer of the corresponding ORF (Harvard Institute of Proteomics) from pDONR201/221 into pAG306GPD-ccdB chr 1 (Hughes and Gottschling, 2012) using Gateway™ LR Clonase™ II Enzyme mix (ThermoFisher) according to the manufacturer’s instructions, followed by sequencing to verify correct integrations. To integrate the resulting expression plasmid into yeast chr I (199456-199457), pAG306GPD-ORF chr 1 was digested with NotI-HF (NEB, #10133990).

### RNA isolation and RT-qPCR

Yeast RNA was isolated and the amount was determined by RT-qPCR as previously described (Schuler et al., 2021) with slight modifications. For RNA isolation, 5 × 10^7^ of overnight log-phase yeast cell cultures grown in high amino acid media were harvested and resuspended in Trizol reagent to a final density of 6 × 10^6^ cells/ml. Cells were lysed with glass beads using an Omni Bead Ruptor 12 Homogenizer. Lysis was performed in six cycles of 20 s and was followed by centrifugation at 10,000 × *g* for one min at 4°C. Supernatants were transferred to a new tube and ethanol was added to a final concentration of 50%. RNA was isolated following the instruction of RNeasy Mini Kit (QIAGEN) and the concentration was determined by Nanodrop. RNA samples were treated with TURBO DNase kit (Invitrogen, AM2238) to clear DNA contamination. High Capacity cDNA Reverse Transcription Kits (Applied Biosystems, 4368814) were used for reverse transcription of RNA samples, followed by RT-qPCR using TaqMan Real-Time PCR Assays (Thermo). MRL1 was used as internal control. Oligo pairs used for RT-qPCR are listed in Table S4.

### Yeast MDC assays

For yeast MDC assays, overnight log-phase cell cultures were directly harvested, or harvested after grown in the presence of DMSO or the indicated drug for 2 h. After incubation, cells were harvested by centrifugation, resuspended in imaging buffer (5% wt/vol Glucose, 10mM HEPES pH 7.6) and optical z-sections of live yeast cells were acquired with an AxioObserver (Carl Zeiss). The percentage of cells with MDCs was quantified from maximum intensity projected wide field images generated in ZEN (Carl Zeiss). Unless otherwise indicated, all quantifications show the mean ± SEM from three biological replicates with n = 100 cells per experiment. MDCs were identified as Tom70-positive Tim50-negative structures of varying size and shape. Maximum intensity projected images are displayed for all yeast images.

### Yeast MDC screen

For the yeast MDC screen, a collection of yeast strains endogenously expressing Tom70-yeGFP and Tim50-mCherry were created by crossing a Tom70-yeGFP Tim50-mCherry query strain (AHY2318) to the yeast non-essential deletion collection (Giaever et al., 2002) using Synthetic Genetic Array (SGA) technology (Tong et al., 2001) with a Biomek Robot (Singer) and standard techniques for high-throughput strain construction. The final library was contained within 96-well plates. The library was grown in YPAD (200 µL for each well) with constant agitation for over 24 h at 30°C to reach stationary phase. The saturated library was then diluted and cultured in YPAD (1 ml for each well) in deep-well 96-well plates with constant agitation for overnight growth at 30°C. In this way, most mutants in the library reach log-phase at a cell density of 2-6 ×10^6^ cells/ml. Strains with low amount of cells for screening were noted in Table S1. The library was centrifuged in deep-well 96-well plates and resuspended in 1ml of YPAD containing 200 nM Rap and cultured for 2 h at 30°C with constant agitation. For imaging, the library was harvested by centrifugation in deep-well 96-well plates, washed once with double-distilled water, and resuspended in 500 µl SD medium supplemented with 2 mg/ml cas-amino acids and 200 nM Rap. The supplementation of amino acids is to ensure consistent MDC formation. The library (100 µl for each well) was then added to glass-bottom 96-well plates pre-coated with 1 mg/ml Concanavalin A to allow the attachment of yeast cells to the bottom of wells. The surfaces of glass-bottom plates contacting water immersion objective were coated with Sigmacote. Optical z-sections of the library were acquired with an AxioObserver (Carl Zeiss). The percentage of cells with MDCs was quantified from widefield images generated in ZEN (Carl Zeiss). At least n=100 cells were assessed, unless the total number of cells was less than 100 as noted in Table S1. Strains with a total number of cells fewer than 5 per well were noted as “low growth” in Fig. 1 B. MDCs were identified as Tom70-enriched, Tim50-negative structures of varying size and shape. Genetic ontology analysis was performed on gene deletions that lead to MDC formation in less than 20%, or higher than 60% of cells using FunSpec (Robinson et al., 2002).

### Microscopy and image analysis

Optical z-sections of live yeast cells were acquired with a Zeiss AxioObserver equipped with a Zeiss Axiocam 506 monochromatic camera, 63× oil-immersion objective (Plan-Apo, NA 1.4). Optical z-sections of live yeast cell libraries for the MDC screen were acquired with a Zeiss AxioObserver equipped with a Zeiss Axiocam 506 monochromatic camera, 63× water-immersion objective (Plan-Apo, NA 1.4). Widefield images were acquired with ZEN (Carl Zeiss). Individual channels of all images were minimally adjusted in Fiji (Schindelin et al., 2012) to match the fluorescence intensities between channels for better visualization.

### Extraction of lipids from yeast whole cell lysates

For analysis of whole cell lysate lipid levels, cells were grown exponentially in the indicated media for 15 h at 30°C to a density of 6-8 ×10^6^ cells/ml and harvested, or harvested after treated with the indicated chemicals for 2 h. 5 × 10^7^ total yeast cells were harvested by centrifugation, washed twice with double-distilled water, and cell pellets were shock frozen in liquid nitrogen.

Extraction of lipids was carried out using a biphasic solvent system of cold methanol, methyl tert-butyl ether (MTBE), and water as described (Matyash et al., 2008) with some modifications. In a randomized sequence, yeast lipids were extracted in bead-mill tubes (glass 0.5 mm, Qiagen, Hilden, Germany) containing a solution of 230 µl MeOH containing internal standards (Avanti SPLASH LipidoMix, Cholesterol-d7 (75 µg/ml), and FA 16:0-d31 (28.8 µg/ml) all at 10 µl per sample) and 250 µl ammonium bicarbonate. Samples were homogenized in one 30 s cycle, transferred to microcentrifuge tubes (polypropylene 1.7 ml, VWR, USA) containing 750 µl MTBE, and rested on ice for 1 h with occasional vortexing. Samples were then centrifuged at 15,000 × *g* for 10 min at 4°C, and the upper phases were collected. A 1 ml aliquot of the upper phase of MTBE/MeOH/water (10:3:2.5, vol/vol/vol) was added to the bottom aqueous layer followed by a brief vortex. Samples were then centrifuged at 15,000 × *g* for 10 min at 4°C, the upper phases were combined and evaporated to dryness under speedvac. Lipid extracts were reconstituted in 500 µl of mobile phase B and transferred to an Liquid chromatography–mass spectrometry (LC–MS) vial for analysis. Concurrently, a process blank sample was prepared and then a pooled quality control (QC) sample was prepared by taking equal volumes (∼50 µl) from each sample after final resuspension.

### Extraction of lipids from isolated yeast mitochondria

Yeast mitochondria were isolated as previously described (Schuler et al., 2016) with some modification. Yeast cells were grown overnight in log-phase as described above, then treated with either DMSO or indicated drugs for 2 h to a density of 6-8 × 10^7^ cells/ml. Cells were then re-isolated by centrifugation, washed with distilled water, and harvested. The pellet weight was determined. Subsequently, cells were resuspended in dithiothreitol (DTT) buffer (0.1 M Tris, 10 mM DTT) for 2 ml per gram of pellets, and incubated for 10 min at 30°C under constant rotation. After re-isolation by centrifugation at 2, 000 × *g*, DTT treated cells were washed once in 1.2 M sorbitol and resuspended in sorbitol phosphate buffer (1.2 M sorbitol, 20 mM K2HPO4, pH = 7.4 with HCl) for 2 ml per gram of pellets. Cell walls were digested in 6.67 ml of sorbitol phosphate buffer per gram of pellet containing 2 mg lyticase per gram of pellets (4mg lyticase per gram of pellets was used when cells were cultured in no amino acid media to allow complete digestion) for 50 min at 30°C under constant rotation. After lyticase digestion, spheroplasts were isolated by centrifugation at 1200 × *g* and lysed by mechanical disruption in 13.3 ml of homogenization buffer (0.6 M sorbitol, 10 mM Tris, pH = 7.4, 1 mM EDTA, pH = 8.0 with KOH, 0.2% BSA, 1 mM PMSF) per gram of pellet at 4°C. Cell debris was removed from the homogenate by centrifugation twice at 1500 × *g* and 2000 × *g* successively for 5 min at 4°C. Crude mitochondria were pelleted at 12000 × *g* for 12 min at 4°C. The mitochondrial pellet was resuspended in 10 ml of SEM buffer (250 mM sucrose, 1 mM EDTA, pH = 8.0 with KOH, 10 mM 3-(N-morpholino)-propansulfonic acid, pH = 7.2), re-isolated by differential centrifugation as described above, and resuspended in SEM buffer. The concentration of mitochondria prepared was determined by Bicinchoninic Acid Protein Assay. For each sample used for lipidomics analysis, 200 µg of mitochondria (by protein) were pelleted at 17500 × *g* for 12 min at 4°C before shock frozen in liquid nitrogen and stored at - 80°C.

Extraction of lipids was carried out using a biphasic solvent system of cold methanol, methyl tert-butyl ether (MTBE), and PBS/water (Matyash et al., 2008) with some modifications. In a randomized sequence, 225 µl MeOH with internal standards described above and 750 µl MTBE was added to each sample. Samples were sonicated for 60 s and then incubated on ice with occasional vortexing for one hour. 188 µl of PBS was then added to induce phase separation and briefly vortexed. Samples were rested at room temperature for 15 min, and then centrifuged at 15,000 × *g* for 10 min at 4°C. The organic (upper) layer was collected, and the aqueous (lower) layer was re-extracted with 1 ml of 10:3:2.5 (vol/vol/vol) MTBE/MeOH/dd-H2O, briefly vortexed, incubated at room temperature, and centrifuged at 15,000 × *g* for 10 min at 4°C. Upper phases were combined and evaporated to dryness under speedvac. Lipid extracts were reconstituted in 450 µL of 4:1:1 (vol/vol/vol) IPA/ACN/water and transferred to an LC-MS vial for analysis. Concurrently, a process blank sample was prepared, and pooled QC samples were prepared by taking equal volumes from each sample after final resuspension.

### LC-MS analysis (QTOF)

Lipid extracts were separated on an Acquity UPLC CSH C18 column (2.1 × 100 mm; 1.7 µm) coupled to an Acquity UPLC CSH C18 VanGuard precolumn (5 × 2.1 mm; 1.7 µm) (Waters, Milford, MA) maintained at 65°C connected to an Agilent HiP 1290 Sampler, Agilent 1290 Infinity pump, and Agilent 6545 Accurate Mass Q-TOF dual AJS-ESI mass spectrometer (Agilent Technologies, Santa Clara, CA). Samples were analyzed in a randomized order in both positive and negative ionization modes in separate experiments acquiring with the scan range m/z 100 - 1700. For positive mode, the source gas temperature was set to 225°C, with a drying gas flow of 11 liters/min, nebulizer pressure of 40 psig, sheath gas temp of 350°C and sheath gas flow of 11 liters/min. VCap voltage is set at 3500 V, nozzle voltage 500V, fragmentor at 110 V, skimmer at 85 V and octopole RF peak at 750 V. For negative mode, the source gas temperature was set to 300°C, with a drying gas flow of 11 liters/min, a nebulizer pressure of 30 psig, sheath gas temp of 350°C and sheath gas flow 11 liters/min. VCap voltage was set at 3500 V, nozzle voltage 75 V, fragmentor at 175 V, skimmer at 75 V and octopole RF peak at 750 V. Mobile phase A consisted of ACN:H2O (60:40, vol/vol) in 10 mM ammonium formate and 0.1% formic acid, and mobile phase B consisted of IPA:ACN:H2O (90:9:1, vol/vol/vol) in 10 mM ammonium formate and 0.1% formic acid. For negative mode analysis the modifiers were changed to 10 mM ammonium acetate. The chromatography gradient for both positive and negative modes started at 15% mobile phase B then increased to 30% B over 2.4 min, it then increased to 48% B from 2.4 - 3.0 min, then increased to 82% B from 3 - 13.2 min, then increased to 99% B from 13.2 - 13.8 min where it is held until 16.7 min and then returned to the initial conditions and equilibrated for 5 min. Flow was 0.4 ml/min throughout, with injection volumes of 5 µl for positive and 10 µl negative mode. Tandem mass spectrometry was conducted using iterative exclusion, the same LC gradient at collision energies of 20 V and 27.5 V in positive and negative modes, respectively.

### LC-MS data processing

For data processing, Agilent MassHunter (MH) Workstation and software packages MH Qualitiative and MH Quantitative were used. The pooled QC (n=8) and process blank (n=4) were injected throughout the sample queue to ensure the reliability of acquired lipidomics data. For lipid annotation, accurate mass and MS/MS matching was used with the Agilent Lipid Annotator library and LipidMatch (Koelmel et al., 2017). Results from the positive and negative ionization modes from Lipid Annotator were merged based on the class of lipid identified. Data exported from MH Quantitative was evaluated using Excel where initial lipid targets are parsed based on the following criteria. Only lipids with relative standard deviations (RSD) less than 30% in QC samples are used for data analysis. Additionally, only lipids with background AUC counts in process blanks that are less than 30% of QC are used for data analysis. The parsed Excel data tables are normalized based on the ratio to class-specific internal standards.

### Quantification and statistical analysis

The number of replicates, what n represents, and dispersion and precision measures are indicated in the figure legends. All experiments were repeated at least three times. All attempts at replication were successful. Sample sizes were as large as possible to be representative, but of a manageable size for quantifications. Specifically, for yeast MDC assays, N = three replicates, with n = 100 cells for each replicate. For lipidomic analysis, N = three to four biological replicates analyzed in the same LC-MS run. All statistical analysis was performed in Prism (GraphPad) and the statistical test used is indicated in the corresponding figure legend. No data were excluded from the analyses, with the exception of some lipidomic samples that did not meet the QC cutoff. In the latter case, all samples of the affected biological replicate were excluded from any further analysis. No randomization or blinding was used as all experiments were performed with defined laboratory reagents and yeast strains of known genotypes.

## Online supplemental material

Fig. S1 shows the lipid profiles of *ups1*Δ, *ups2*Δ, *ups1Δups2*Δ, *mdm35*Δ, *gep4*Δ, and *crd1*Δ yeast strains, and quantifies the percentage of the indicated mutant cells forming MDCs under the indicated MDC-inducing conditions. Fig. S2 shows the lipid profiles of *psd1*Δ, *psd2*Δ, and *psd1Δpsd2*Δ yeast strains, the mRNA level of *CHO1* in the indicated mutant cells in response to the indicated treatments and quantifies the percentage of the indicated mutant cells forming MDCs under the indicated MDC-inducing conditions. Fig. S3 shows the PE level changes in the indicated mutant cells in response to the indicated treatments, and demonstrates that overexpression of *UPS2* and *PSD1* affect MDC biogenesis and increase cellular MMPE and DMPE abundance. Fig. S4 shows the dependency of MDC biogenesis and PE decline on amino acid abundance, deletion of *PSD1* bypasses amino acid stress and triggers MDC formation, and demonstrates that *PAH1* deletion does not affect MDC formation. Table S1 lists the quantification results of the MDC screen and gene ontology analysis. Table S2 lists the bacterial strains, chemicals, plasmids, and software used in this study. Table S3 lists the yeast strains used in this study. Table S4 lists the oligonucleotides used in this study.

## ACKNOWLEDGEMENTS

We thank members of Adam L. Hughes Laboratory for discussion and manuscript comments. We thank members of Janet. M. Shaw Laboratory for technical assistance. We thank Jenna M. Goodrum for helping construct the yeast deletion collection used for the preliminary screen. We thank Janet M. Shaw for contributing stipend support for T. Xiao. We thank Dan Cuthbertson of Agilent Technologies for assistant with tandem mass spectrometry.

Metabolomics analysis was performed at the Metabolomics Core Facility at the University of Utah. Mass spectrometry equipment was obtained through the NCRR Shared Instrumentation Grant 1S10OD016232-01, 1S10OD018210-01A1 and 1S10OD021505-01. Research was supported by National Institutes of Health grant GM119694 (to A.L.H.); NIH T32GM007464 (to A.M.E.); a University of Utah Graduate Research Fellowship (to T.X.); and the Howard Hughes Medical Institute (to J.M.S.).

## Author contributions

T. Xiao. and A.L. Hughes. conceived the study; T. Xiao, A.M. English, and J. Alan M. performed and analyzed experiments; T. Xiao and A.M. English performed the preliminary genetic screen; J. Alan M. performed lipidomic analysis; J.E. Cox supervised the lipidomic analysis; T. Xiao and A.L. Hughes wrote the manuscript; T. Xiao, A.M. English, J. Alan M., and A.L. Hughes edited the manuscript; A.L. Hughes supervised the study.

## SUPPLEMENTAL FIGURE and TABLE LEGENDS

**Figure S1.**
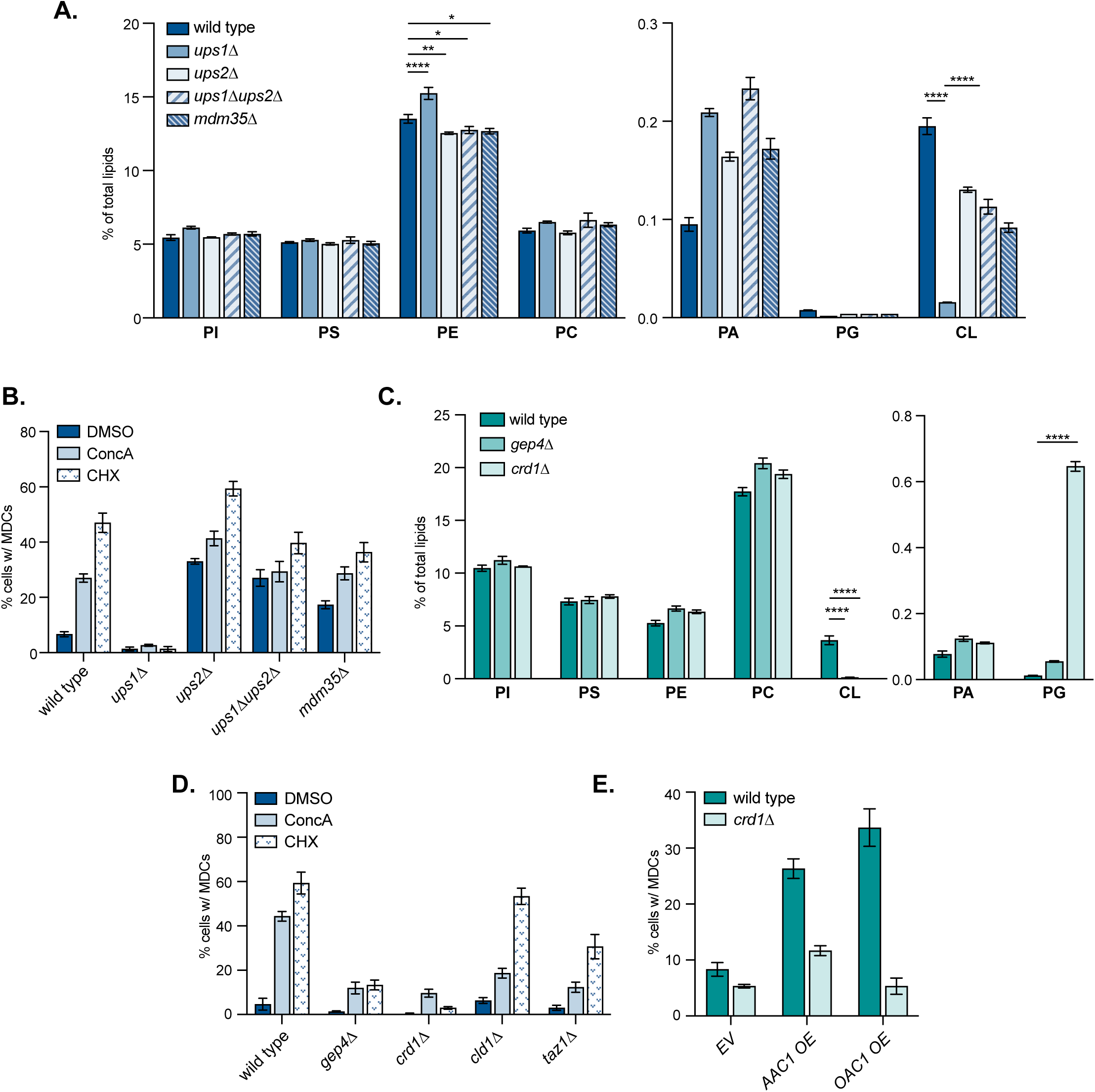
Suppressing CL production inhibits MDC formation (related to Figures 2-3). **(A)** The relative amounts of the indicated phospholipids in whole-cell lysates of wild-type or the indicated mutant cells determined by mass spectrometry-based lipidomic analysis. Amount of each lipid relative to total lipids were determined. Error bars = SEM of three replicates. Statistical comparison shows difference to the corresponding wild-type control. n.s., not significant, *p<0.0332, **p<0.0021, ***p<0.0002, ****p<0.0001, 2-way ANOVA with Holm-ŠÍdák test. **(B)** Quantification of MDC formation in wild-type cells or the indicated mutant cells treated with DMSO, ConcA, or CHX for 2 h. N > 100 cells per replicate, error bars = SEM of three replicates. **(C)** The relative amounts of the indicated phospholipids in whole-cell lysates of wild-type or the indicated mutant cells determined by mass spectrometry-based lipidomic analysis. Amounts of each lipid relative to total lipids were determined. Error bars = SEM of three replicates. Statistical comparison shows difference to the corresponding wild-type control. n.s., not significant, *p<0.0332, **p<0.0021, ***p<0.0002, ****p<0.0001, 2-way ANOVA with Holm-ŠÍdák test. **(D)** Quantification of MDC formation in wild-type cells or the indicated mutant cells treated with DMSO, ConcA, or CHX for 2 h. N > 100 cells per replicate, error bars = SEM of three replicates. **(E)** Quantification of MDC formation in wild-type cells or *crd1*Δ cells integrated with empty vector (*EV*), *AAC1* overexpressing vector (*AAC1 OE*), or *OAC1* overexpressing vector (*OAC1 OE*). N > 100 cells per replicate, error bars = SEM of three replicates.

**Figure S2.**
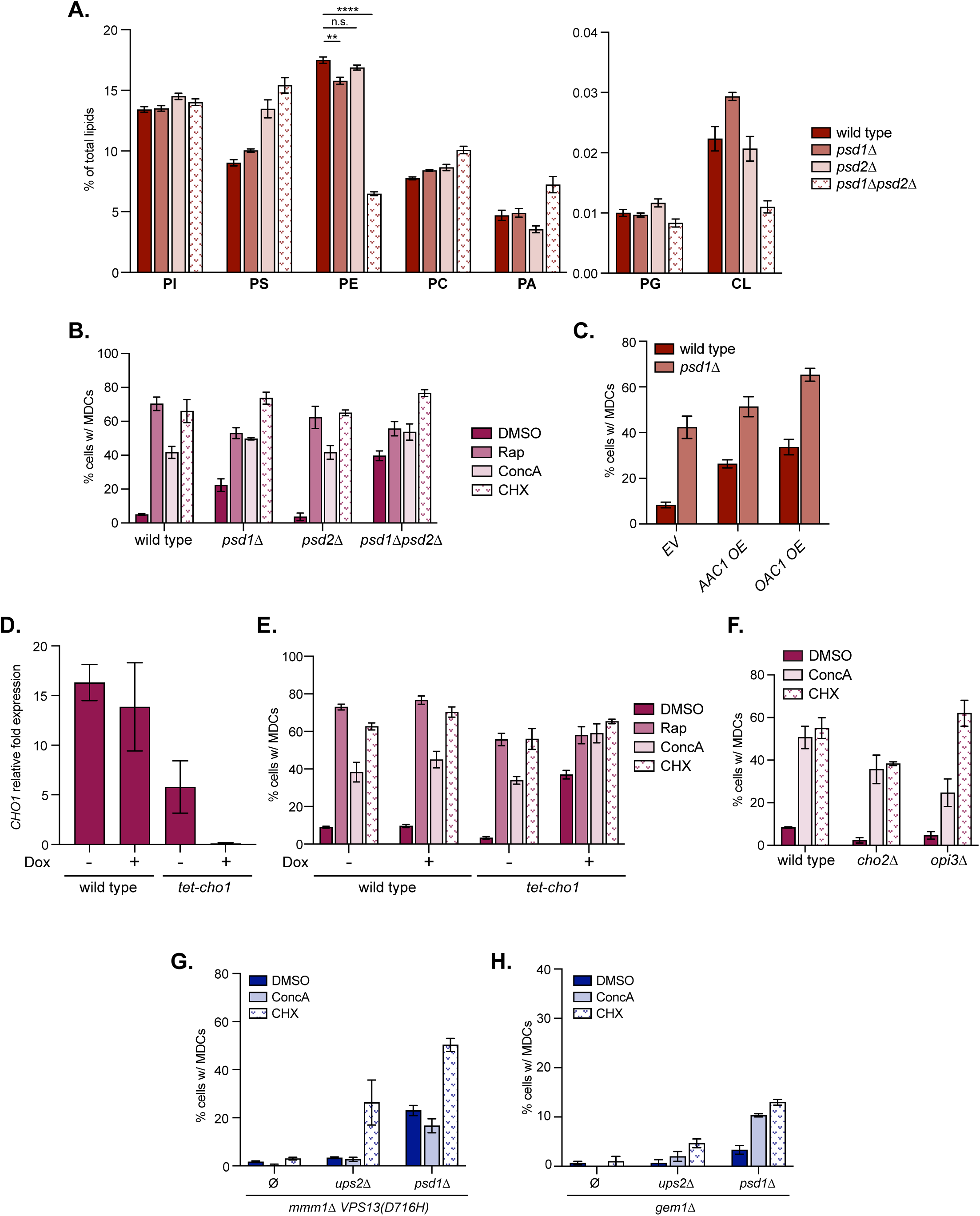
Inhibiting mitochondrial PE synthesis activates MDC biogenesis (related to Figures 4-5). **(A)** The relative amounts of the indicated phospholipids in whole-cell lysates of wild-type or the indicated mutant cells determined by mass spectrometry-based lipidomic analysis. Amounts of each lipid relative to total lipids were determined. Error bars = SEM of three replicates. Statistical comparison shows difference to the corresponding wild-type control. n.s., not significant, *p<0.0332, **p<0.0021, ***p<0.0002, ****p<0.0001, 2-way ANOVA with Holm-ŠÍdák test. **(B)** Quantification of MDC formation in wild-type cells or the indicated mutant cells treated with DMSO, Rap, ConcA, or CHX for 2 h. N > 100 cells per replicate, error bars = SEM of three replicates. **(C)** Quantification of MDC formation in wild-type cells or *psd1*Δ cells with genomic integration of an empty vector (*EV*), *AAC1* overexpressing vector (*AAC1 OE*), or *OAC1* overexpressing vector (*OAC1 OE*). N > 100 cells per replicate, error bars = SEM of three replicates. **(D)** Quantification of *CHO1* mRNA abundance by RT-qPCR in wild-type cells or *tet-cho1* mutant cells treated with DMSO, Rap, ConcA, or CHX for 2 h in the absence or presence of Dox. Error bars = SEM of three replicates **(E)** Quantification of MDC formation in wild-type cells or *tet-cho1* cells treated with DMSO, Rap, ConcA, or CHX for 2 h in the absence or presence of Dox. N > 100 cells per replicate, error bars = SEM of three replicates. **(F)** Quantification of MDC formation in wild-type cells or the indicated mutant cells treated with DMSO, Rap, ConcA, or CHX for 2 h. N > 100 cells per replicate, error bars = SEM of three replicates. **(G)**Quantification of MDC formation in *mmm1Δ VPS13(D716H)* cells or *mmm1Δ VPS13(D716H)* cells with the indicated gene deletions treated with DMSO, ConcA, or CHX for 2 h. Ø, no gene is deleted. N > 100 cells per replicate, error bars = SEM of three replicates. **(H)** Quantification of MDC formation in *gem1*Δ mutant cells or *gem1*Δ cells with the indicated gene deletions treated with DMSO, ConcA, or CHX for 2 h. Ø, no gene is deleted. N > 100 cells per replicate, error bars = SEM of three replicates.

**Figure S3.**
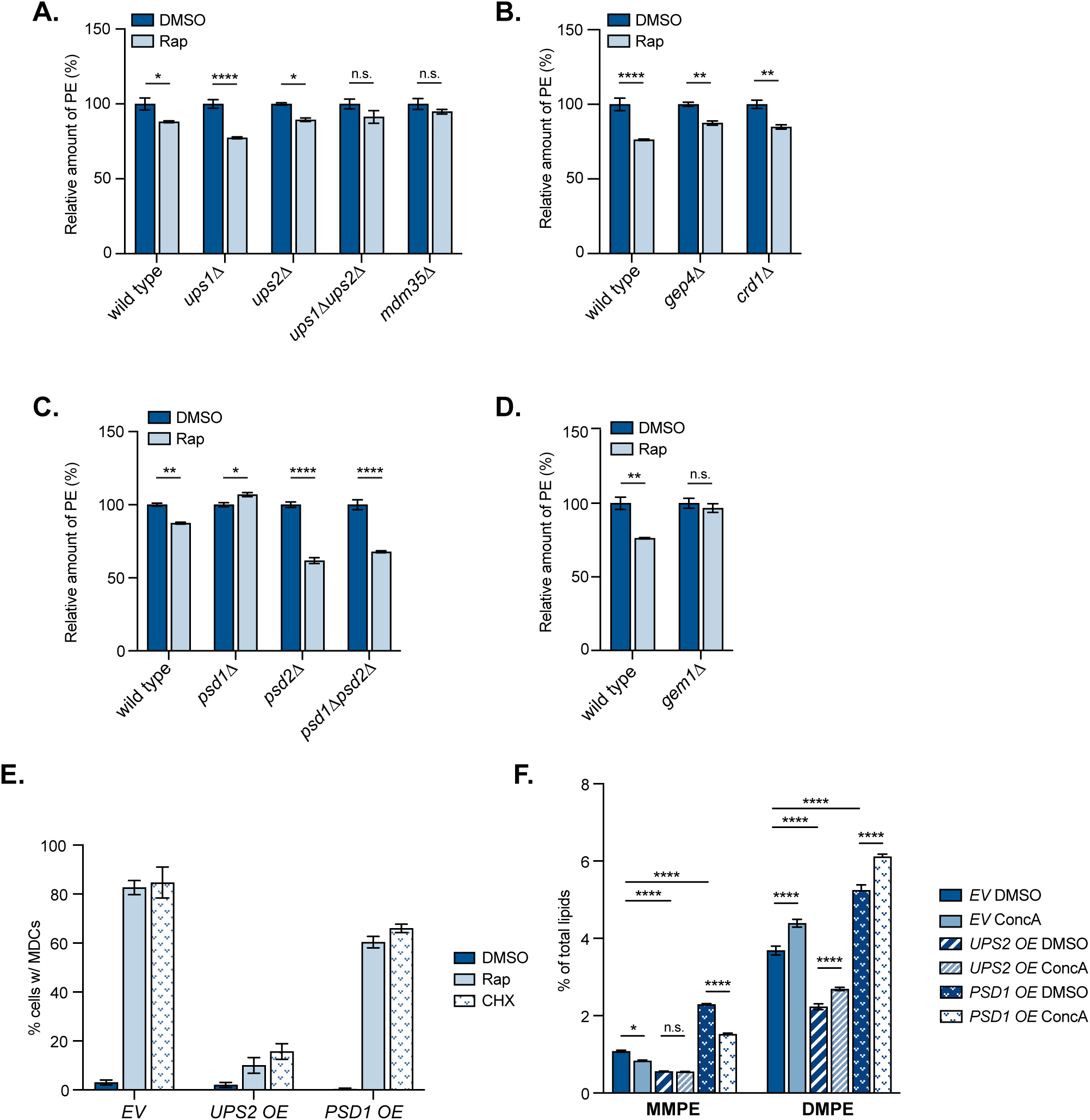
MDC-inducing conditions trigger PE depletion (related to Figure 6). **(A-D)** The relative amounts of PE in whole-cell lysate of wild-type cells or the indicated mutant yeast cells treated with DMSO or Rap for 2 h. The amounts of PE in DMSO-treated cells were set to 100%. Error bars = SEM of three replicates. Statistical comparison shows difference to the corresponding DMSO control. n.s., not significant, *p<0.0332, **p<0.0021, ***p<0.0002, ****p<0.0001, 2-way ANOVA with Holm-ŠÍdák test. **(E)** Quantification of MDC formation in cells with genomic integration of an empty vector (*EV*), *UPS2* overexpressing vector (*UPS2 OE*), or *PSD1* overexpressing vector (*PSD1 OE*) treated with DMSO, Rap, or CHX for 2 h. N > 100 cells per replicate, error bars = SEM of three replicates. **(F)** The relative amounts of MMPE and DMPE in whole-cell lysate of yeast cells with genomic integration of an empty vector (*EV*), *UPS2* overexpressing vector (*UPS2 OE*), or *PSD1* overexpressing vector (*PSD1 OE*) treated with DMSO or ConcA for 2 h. Amounts of each lipid relative to total lipids were determined. Error bars = SEM of three replicates. Statistical comparison shows difference to the corresponding DMSO control. n.s., not significant, *p<0.0332, **p<0.0021, ***p<0.0002, ****p<0.0001, 2-way ANOVA with Holm-ŠÍdák test.

**Figure S4.**
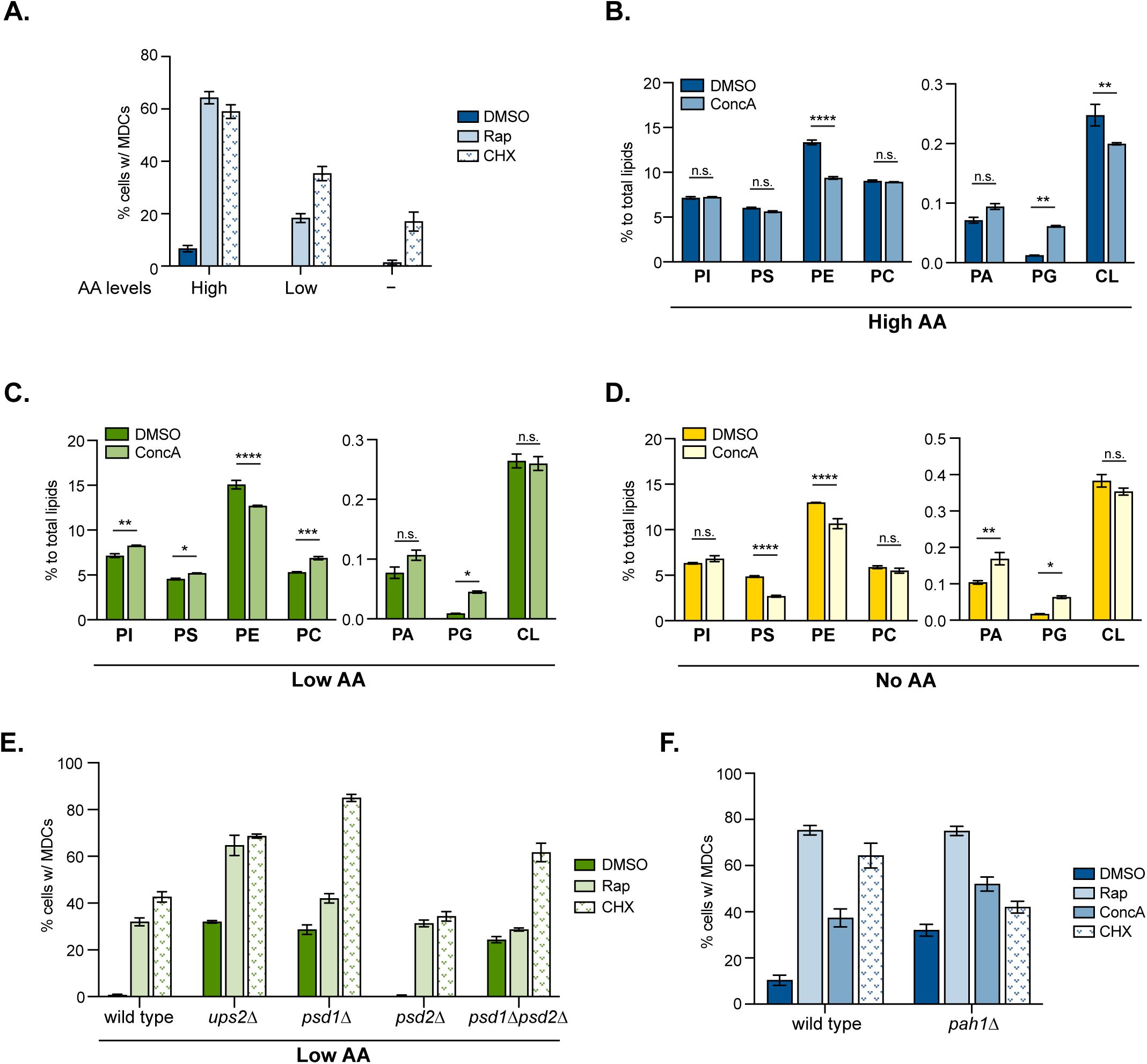
PE decline activates MDC formation under amino acid stress (related to Figures 7-8). **(A)** Quantification of MDC formation in yeast cells grown in amino acid-rich media, synthetic media that contains low levels of amino acid, or minimal media that excludes amino acids (indicated as High, Low, and None, respectively) treated with DMSO, Rap, or CHX for 2 h. N > 100 cells per replicate, error bars = SEM of three replicates. **(B-D)** The relative amounts of the indicated phospholipids in whole-cell lysates of yeast cells grown in amino acid-rich media (High AA) (B), synthetic media that contains low levels of amino acid (Low AA) (C), or minimal media that excludes amino acids (No AA) (D) treated with DMSO or ConcA for 2 h detected by mass spectrometry-based lipidomic analysis. Amounts of each lipid relative to total lipids were determined. Error bars = SEM of three replicates. Statistical comparison shows difference to the corresponding DMSO control. n.s., not significant, *p<0.0332, **p<0.0021, ***p<0.0002, ****p<0.0001, 2-way ANOVA with Holm-ŠÍdák test. **(E)** Quantification of MDC formation in wild-type cells or the indicated mutant cells grown in synthetic media that contains low levels of amino acid (Low AA) treated with DMSO, Rap, or CHX for 2 h. N > 100 cells per replicate, error bars = SEM of three replicates. **(F)** Quantification of MDC formation in wild-type cells or *pah1*Δ cells treated with DMSO, Rap, ConcA, or CHX for 2 h. N > 100 cells per replicate, error bars = SEM of three replicates.

**Table S1. Preliminary screen results of MDC formation in yeast deletion collection mutants.**

**(Tab 1)** Raw results recording the plate number, positions in 96-well plates, cell growth, quantification of cells forming MDCs, % of cells that form MDCs, and notes about cell/mitochondrial morphology of individual mutants of yeast non-essential deletion collections.

**(Tab 2)** Calculation of the ratio of mutants that could not screened (no growth or low image quality), grew poorly (≤ 5 cells per well), or formed MDCs in the indicated percentages of cells, to the total number of open reading frames (ORFs) contained in the yeast deletion collection.

**(Tab 3)** Summary of mutants that show reduced (<20% cells form MDCs in response to Rap) or enhanced (<60% cells form MDCs in response to Rap) MDC formation. Mutants the form MDCs in 0-10%, 10-20%, 60-70%, and >70% of cells are indicated by dark blue, light blue, yellow, and brown, respectively.

**(Tab 4)** Summary of genetic ontology analysis performed with mutants in Tab3.

**Table S2. Bacterial strains, chemicals, plasmids, and software used in this study**

**Table S3. Yeast strains used in this study.**

**Table S4. Oligos used in this study.**

