## Supplemental Table 2 for "Genetic Screen Uncovers a Dual Role for Phospholipids in Mitochondrial-Derived Compartment Biogenesis"

**Table S2. Bacterial strains, chemicals, plasmids, and software used in this study**

| Reagent or resource | Source | Identifier |
|---|---|---|
| **Bacterial strains** | | |
| *Escherichia coli* DH5α | N/A | N/A |
| *S. cerevisiae* ORF collection (pDONR201/221) | Harvard Institute of Proteomics | N/A |
| One Shot™ *ccd*B Survival™ 2 T1$^{R}$ Competent Cells | Thermo Fisher | Cat#A10460 |
| **Chemicals, peptides, and recombinant proteins** | | |
| Casamino acids | US Biological | Cat # 0012501A; CAS # 65072-00-6 |
| Concanamycin A | Santa Cruz Biotechnology | Cat # sc-202111; CAS # 80890-47-7 |
| Concanavalin A | Sigma-Aldrich | Cat # L7647; CAS # 11028-71-0 |
| Cycloheximide | Sigma-Aldrich | Cat # C1988; CAS # 66-81-9 |
| Dimethyl sulfoxide (DMSO) | Sigma-Aldrich | Cat # D2650; CAS # 67-68-5 |
| Doxycycline hyclate | Sigma-Aldrich | Cat # D9891; CAS # 24390-14-5 |
| DTT | Gold Biotechnology | Cat # DTT10; CAS # 27565-41-9 / 3483-12-3 |
| Lyticase | Sigma-Aldrich | Cat # L2524; CAS # 37340-57-1 |
| Rapamycin | LC Laboratories | Cat # R-5000; CAS # 53123-88-9 |
| Sigmacote® | Sigma-Aldrich | Cat # SL2 |
| Critical commercial assays | | |
| Bicinchoninic Acid Protein Assay | Thermo Fisher | Cat # 23227 |
| Gateway LR Clonase II Enzyme Mix | Thermo Fisher | Cat # 11791020 |
| **Plasmids** | | |
| Plasmid: pAG306GPD-AAC1 chr 1 | Schuler et al., 2021 | N/A |
| Plasmid: pAG306GPD-ccdB chr 1 | Hughes and Gottschling, 2012 | N/A |
| Plasmid: pAG306GPD-OAC1 chr 1 | Schuler et al. 2021 | N/A |
| Plasmid: pAG306GPD-PSD1 chr 1 | This study | N/A |

| | | |
|---|---|---|
| Plasmid: pAG306GPD-UPS2 chr 1 | This study | N/A |
| Plasmid: pHLUM | Mülleder *et al*., 2012; Addgene | Plasmid # 40276 |
| Plasmid: pKT127-mCherry | Daniel Gottschling (Calico) | N/A |
| Plasmid: pKT128 | Sheff and Thorn, 2004; Addgene | Plasmid # 8729 |
| Plasmid: pRS306 | Sikorski and Hieter, 1989 | N/A |
| Plasmid: pRS40Hyg | Daniel Gottschling (Calico) | N/A |
| Plasmid: pRS40Nat | Daniel Gottschling (Calico) | N/A |
| **Software and algorithms** | | |
| FIJI | Schindelin *et al*., 2012 | Version 1 |
| FunSpec | Robinson *et al*., 2002 | http://funspec.med.utoronto.ca |
| LipidMatch | Koelmel *et al*., 2017 | N/A |
| Prism | GraphPad Software, Inc. | Version 9 |
| SnapGene | GSL Biotech | Version 4.2 |
| ZEN Blue Edition | Carl Zeiss Microscopy | Version 2.6 |
