## Supplemental Table 3 for "Genetic Screen Uncovers a Dual Role for Phospholipids in Mitochondrial-Derived Compartment Biogenesis"

| Table S3. Yeast strains used in this study. | |
| --- | --- |
| **Genotype** | **Strain** |
| BY4741 MATa his3Δ1 leu2Δ0 ura3Δ0 met15Δ0 | |
| BY4742 MATα his3Δ1 leu2Δ0 ura3Δ0 lys2Δ0 | |
| BY4743 MATa/MATα his3Δ1/his3Δ1 leu2Δ0/leu2Δ0 ura3Δ0/ura3Δ0 met15Δ0/+ lys2Δ0/+ | |
| MATα TOM70-yEGFP:NatMX TIM50-mCherry:HygMX can1Δ::STE2p-Sp_HIS5 lyp1Δ his3Δ1 leu2Δ0 ura3Δ0 met15Δ0 | AHY 2318 |
| MATa URA3::CMV-tTA his3Δ1 leu2Δ0 ura3Δ0 met15Δ0 | AHY 2677 |
| BY4741 gem1Δ::HygMX | AHY 4230 |
| MATa mmm1Δ::HygMX TOM70-yEGFP:KanMX TIM50-mCherry:KanMX his3Δ1 leu2Δ0 ura3Δ0 lys2Δ0 pVPS13(D716H) | AHY 6760 |
| BY4741 TOM70-yEGFP:HisMX TIM50-mCherry:KanMX | AHY 7019 |
| BY4742 TOM70-yEGFP:HisMX TIM50-mCherry:KanMX | AHY 7021 |
| MATa URA3::CMV-tTA TOM70-GFP:spHis5MX TIM50-mCherry:KanMX his3Δ1 leu2Δ0 ura3Δ0 met15Δ0 | AHY 9543 |
| BY4743 TOM70-yEGFP:KanMX/+ TIM50-mCherry:KanMX/+ chr I(199456-199457)::$P_{GPD1}$-Term$_{CYC1}$-URA3/+ | AHY 10568 |
| BY4743 TOM70-yEGFP:KanMX/+ TIM50-mCherry:KanMX/+ chr I(199456-199457)::$P_{GPD1}$-UPS2-Term$_{CYC1}$-URA3/+ | AHY 11351 |
| BY4743 TOM70-yEGFP:KanMX/+ TIM50-mCherry:KanMX/+ chr I(199456-199457)::$P_{GPD1}$-PSD1-Term$_{CYC1}$-URA3/+ | AHY 11353 |
| MATa psd1Δ::NatMX gem1Δ::HygMX TOM70-yEGFP:HisMX TIM50-mCherry:KanMX his3Δ1 leu2Δ0 ura3Δ0 lys2Δ0 | AHY 11357 |
| MATa psd1Δ::NatMX TOM70-yEGFP:HisMX TIM50-mCherry:KanMX his3Δ1 leu2Δ0 ura3Δ0 lys2Δ0 | AHY 11359 |
| MATa psd1Δ::NatMX his3Δ1 leu2Δ0 ura3Δ0 | AHY 11362 |
| MATα mdm35Δ::NatMX TOM70-yEGFP:HisMX TIM50-mCherry:KanMX his3Δ1 leu2Δ0 ura3Δ0 | AHY 11367 |
| MATa mdm35Δ::NatMX his3Δ1 leu2Δ0 ura3Δ0 lys2Δ0 | AHY 11370 |
| MATa ups2Δ::NatMX gem1Δ::HygMX TOM70-yEGFP:HisMX TIM50-mCherry:KanMX his3Δ1 leu2Δ0 ura3Δ0 met15Δ0 | AHY 11435 |
| MATα ups2Δ::NatMX TOM70-yEGFP:HisMX TIM50-mCherry:KanMX his3Δ1 leu2Δ0 ura3Δ0 lys2Δ0 | AHY 11438 |
| MATa ups2Δ::NatMX his3Δ1 leu2Δ0 ura3Δ0 met15Δ0 | AHY 11440 |
| MATa gem1Δ::HygMX TOM70-yEGFP:HisMX TIM50-mCherry:KanMX his3Δ1 leu2Δ0 ura3Δ0 | AHY 11442 |
| BY4741 gep4Δ::URA3 TOM70-yEGFP:KanMX TIM50-mCherry:KanMX | AHY 11533 |
| MATa ups1Δ::HygMX ups2Δ::NatMX TOM70-yEGFP:HisMX TIM50-mCherry:KanMX his3Δ1 leu2Δ0 ura3Δ0 lys2Δ0 | AHY 11634 |
| MATa ups1Δ::HygMX ups2Δ::NatMX his3Δ1 leu2Δ0 ura3Δ0 met15Δ0 | AHY 11636 |

| | |
|---|---|
| MATa opi3Δ::URA3 TOM70-yEGFP:HisMX TIM50-mCherry:KanMX his3Δ1 leu2Δ0 ura3Δ0 | AHY 11766 |
| MATa cho2Δ::URA3 TOM70-yEGFP:HisMX TIM50-mCherry:KanMX his3Δ1 leu2Δ0 ura3Δ0 lys2Δ0 met15Δ0 | AHY 11768 |
| MATa URA3::CMV-tTA KanMX-tetO7-TATA-CHO1 his3Δ1 leu2Δ0 ura3Δ0 met15Δ0 | AHY 11833 |
| MATa psd1Δ::NatMX psd2Δ::HygMX his3Δ1 leu2Δ0 ura3Δ0 lys2Δ0 met15Δ0 | AHY 11874 |
| MATa psd2Δ::HygMX his3Δ1 leu2Δ0 ura3Δ0 lys2Δ0 | AHY 11964 |
| MATa psd2Δ::HygMX TOM70-yEGFP:HisMX TIM50-mCherry:KanMX his3Δ1 leu2Δ0 ura3Δ0 | AHY 11966 |
| BY4741 TOM70-yEGFP:KanMX TIM50-mCherry:KanMX pHLUM | AHY 11976 |
| MATa URA3::CMV-tTA KanMX-tetO7-TATA-CHO1 TOM70-yEGFP:HisMX TIM50-mCherry:HygMX his3Δ1 leu2Δ0 ura3Δ0 met15Δ0 | AHY 11990 |
| MATa psd1Δ::NatMX psd2Δ::HygMX TOM70-yEGFP:HisMX TIM50-mCherry:KanMX his3Δ1 leu2Δ0 ura3Δ0 met15Δ0 | AHY 12011 |
| MATa crd1Δ::HygMX his3Δ1 leu2Δ0 ura3Δ0 met15Δ0 | AHY 12516 |
| MATa crd1Δ::NatMX TOM70-yEGFP:HisMX TIM50-mCherry:KanMX his3Δ1 leu2Δ0 ura3Δ0 met15Δ0 | AHY 12518 |
| MATα pah1Δ::NatMX TOM70-yEGFP:HisMX TIM50-mCherry:KanMX his3Δ1 leu2Δ0 ura3Δ0 met15Δ0 | AHY 12555 |
| MATa gep4Δ::HygMX his3Δ1 leu2Δ0 ura3Δ0 met15Δ0 | AHY 12736 |
| MATa taz1Δ::NatMX TOM70-yEGFP:HisMX TIM50-mCherry:KanMX his3Δ1 leu2Δ0 ura3Δ0 met15Δ0 | AHY 12775 |
| MATa ups1Δ::HygMX his3Δ1 leu2Δ0 ura3Δ0 met15Δ0 | AHY 12801 |
| MATα ups1Δ::HygMX TOM70-yEGFP:HisMX TIM50-mCherry:KanMX his3Δ1 leu2Δ0 ura3Δ0 | AHY 12803 |
| BY4741 pHLUM | AHY 12872 |
| MATa/MATα psd1Δ::NatMX/psd1Δ::NatMX TOM70-yEGFP:HisMX/+ TIM50-mCherry:KanMX/+ his3Δ1/his3Δ1 leu2Δ0/leu2Δ0 ura3Δ0/ura3Δ0 lys2Δ0/+ met15Δ0/+ chr I(199456-199457)::$P_{GPD1}$-$Term_{CYC1}$-URA3/+ | AHY 13656 |
| BY 4743 chr I(199456-199457)::$P_{GPD1}$-UPS2-$Term_{CYC1}$-URA3/+ | AHY 13707 |
| BY 4743 chr I(199456-199457)::$P_{GPD1}$-PSD1-$Term_{CYC1}$-URA3/+ | AHY 13709 |
| BY 4743 chr I(199456-199457)::$P_{GPD1}$-$Term_{CYC1}$-URA3/+ | AHY 13713 |
| MATa ups2Δ::URA3 mmm1Δ::HygMX TOM70-yEGFP:KanMX TIM50-mCherry:KanMX his3Δ1 leu2Δ0 ura3Δ0 lys2Δ0 pVPS13(D716H) | AHY 14155 |
| MATa psd1Δ::URA3 mmm1Δ::HygMX TOM70-yEGFP:KanMX TIM50-mCherry:KanMX his3Δ1 leu2Δ0 ura3Δ0 lys2Δ0 pVPS13(D716H) | AHY 14157 |
| MATa/MATα psd1Δ::NatMX/psd1Δ::NatMX TOM70-yEGFP:HisMX/+ TIM50-mCherry:KanMX/+ his3Δ1/his3Δ1 leu2Δ0/leu2Δ0 ura3Δ0/ura3Δ0 lys2Δ0/+ met15Δ0/+ chr I(199456-199457)::$P_{GPD1}$-AAC1-$Term_{CYC1}$-URA3/+ | AHY 14226 |

| | |
|---|---|
| MATa/MATα psd1Δ::NatMX/psd1Δ::NatMX TOM70-yEGFP:HisMX/+ TIM50-mCherry:KanMX/+ his3Δ1/his3Δ1 leu2Δ0/leu2Δ0 ura3Δ0/ura3Δ0 lys2Δ0/+ met15Δ0/+ chr I(199456-199457)::$P_{GPD1}$-OAC1-$Term_{CYC1}$-URA3/+ | AHY 14228 |
| MATa/MATα crd1Δ::NatMX/crd1Δ::NatMX TOM70-yEGFP:HisMX/+ TIM50-mCherry:KanMX/+ his3Δ1/his3Δ1 leu2Δ0/leu2Δ0 ura3Δ0/ura3Δ0 met15Δ0/+ chr I(199456-199457)::$P_{GPD1}$-$Term_{CYC1}$-URA3/+ | AHY 14230 |
| MATa/MATα crd1Δ::NatMX/crd1Δ::NatMX TOM70-yEGFP:HisMX/+ TIM50-mCherry:KanMX/+ his3Δ1/his3Δ1 leu2Δ0/leu2Δ0 ura3Δ0/ura3Δ0 met15Δ0/+ chr I(199456-199457)::$P_{GPD1}$-AAC1-$Term_{CYC1}$-URA3/+ | AHY 14232 |
| MATa/MATα crd1Δ::NatMX/crd1Δ::NatMX TOM70-yEGFP:HisMX/+ TIM50-mCherry:KanMX/+ his3Δ1/his3Δ1 leu2Δ0/leu2Δ0 ura3Δ0/ura3Δ0 met15Δ0/+ chr I(199456-199457)::$P_{GPD1}$-OAC1-$Term_{CYC1}$-URA3/+ | AHY 14234 |
| MATa cld1Δ::NatMX TOM70-yEGFP:HisMX TIM50-mCherry:KanMX his3Δ1 leu2Δ0 ura3Δ0 met15Δ0 | AHY 14261 |
