## Supplemental Table 4 for "Genetic Screen Uncovers a Dual Role for Phospholipids in Mitochondrial-Derived Compartment Biogenesis"

| Table S4. Oligos used in this study. | | |
|---|---|---|
| **Name** | **Sequence** | **Number** |
| AmpR R | CTGATCTTCAGCATCTTTTACTTTCACCAGCG | 2531 |
| CHO2 qPCR F | CACACGGACACAGACGTTAT | 5461 |
| CHO2 qPCR R | TGAGGCCCTTCTGCTTAATG | 5462 |
| CHO2 KO check F | AATTGGTCAAGACAGTCAATTGCCACACTT | 3942 |
| CHO2 KO check R | GGAAGGATGATGGTGACGATGAAG | 4767 |
| CHO2 KO F | AGTGATTTTCTTAGTGACAAAGCTTTTTCTTCATCT GTAGagattgtactgagagtgcac | 3940 |
| CHO2 KO R | CCTAGTACTTTTTAAATATATATACTCAAAAAAAAA AACctgtgcggtatttcacaccg | 3941 |
| CLD1 KO check F | TGCAGTTCGTCATTAATATGTCAGGATGGA | 5404 |
| CLD1 KO check R | CTAACCCCCGAGAAGCATTAAG | 5405 |
| CLD1 KO F | ACACTAATACTTATACTGATTAATAAGGGTTAGCCT TTTAagattgtactgagagtgcac | 5402 |
| CLD1 KO R | AAAATTTCGTTATATAATGTAGATGCACAAGATTTC TTCActgtgcggtatttcacaccg | 5403 |
| CRD1 KO check F | TATTGGAACTCTGAACCTGTAACAACCAGT | 1279 |
| CRD1 KO F | CAGGCCTGGTAGCATAGTTTGGTCCCTAATAATTT AGTCAAGATTGTACTGAGAGTGCAC | 1277 |
| CRD1 KO R | TGAAAAGTCAGGACCCTTTTCAAAAAGGATCGCAA TTATACTGTGCGGTATTTCACACCG | 1278 |
| Gateway Chr I Intg Chk R | GCACTACGTCAATGCAAGT | 3148 |
| GEM1 KO check F | GATGAAGAATTTATCCCAATATTAATGGAG | 3074 |
| GEM1 KO check R | CATAAGTATCTAAACCGAAGATTTCTTCAC | 3236 |
| GEM1 KO F | AAATAGCGGACTTCTAAATACTAATGTGTTGAACAA CACAGATTGTACTGAGAGTGCACC | 577 |
| GEM1 KO R | GAAATGCAACACTTCCCTAATATAGAAATTTGGGC ATTAACTGTGCGGTATTTCACACCG | 578 |
| GEP4 KO check F | TTCAGAGCCACCTTCGGAGTATGTCTCAAT | 4676 |
| GEP4 KO check R | GTCTTCCACCAGATTCTCCG | 4677 |
| GEP4 KO F | AAAGGCGGCAGTTACATTACATCGTCTCCTCTACC TAGTCagattgtactgagagtgcac | 4674 |
| GEP4 KO R | AAAAAATTAAAATGTTTTACTTTTTATTAAAGTTGCC TAActgtgcggtatttcacaccg | 4675 |
| MDM35 KO check F | CTAGTTCCAGCATGGTACCGTCTCCGATTG | 4482 |
| MDM35 KO check R | GTCCAGTCAGGGAACAATTCTG | 4483 |
| MDM35 KO F | GTTTTAACTTGAATTACAATAACAATAATACCAGTTT TATagattgtactgagagtgcac | 4480 |
| MDM35 KO R | ACATGTTGAATAATGCACATTCTGTGCTAAAATATA TACTctgtgcggtatttcacaccg | 4481 |
| MMM1 KO check F | TTGATTGTGCACTCGTAAGTGACTTGACTG | 567 |

| | | |
|---|---|---|
| MMM1 KO check R | GATCAATACACATTGTCAACTATAATGC | 918 |
| MMM1 KO F | TTGAGAGAGTCAATATAATACCTGTAGCCTTTTTCT GAAAGATTGTACTGAGAGTGCACC | 565 |
| MMM1 KO R | GATAGGAAAAAGATAGAACAAAAAATTTGTACATAA ATATCTGTGCGGTATTTCACACCG | 566 |
| MRL1 qPCR F | ggggcataagactacggtca | 4363 |
| MRL1 qPCR R | gaactccctgttggcaagg | 4363 |
| OPI3 KO check F | GTCGACCGTCTCCAAGAGATCCACGATAAT | 4765 |
| OPI3 KO check R | CCTGGAGTGTCGATGATGCAG | 4766 |
| OPI3 KO F | AACAGCAATTGAAGACAACAAGAATAGCGCAAGTC AAGCGagattgtactgagagtgcac | 4763 |
| OPI3 KO R | ATAGGCTTCTAACATTATAGAATATATAGAAATAGA GCACctgtgcggtatttcacaccg | 4764 |
| PAH1 KO check F | GTTCGCAGTTCCTAACACTGAGCGTTCTTG | 4941 |
| PAH1 KO check R | CCTCTTCCTGTTATACACGCATTG | 4942 |
| PAH1 KO F | ACAGGGAAGAAATTACTGAAGATAGACACATCGGT CGATTagattgtactgagagtgcac | 4939 |
| PAH1 KO R | AGTATGGATCGTTATAAATAATATTCGGCTACAAGA ATCTctgtgcggtatttcacaccg | 4940 |
| PSD1 KO check F | TGACCGTGTTCACTGTGAGCTATTGCAGAA | 1276 |
| PSD1 KO check R | GACAATGGTGGAACTCGATACTC | 4678 |
| PSD1 KO F | GGTCGTTATTTTTTGAAGAAGAAGGAAAAGCAAAG CCAGCAGATTGTACTGAGAGTGCAC | 1274 |
| PSD1 KO R | TATACAGCAAAATAAATGCTAACTTTACATATGATT GCTTCTGTGCGGTATTTCACACCG | 1275 |
| PSD2 KO check F | AAC TAC ACT TGC ATT ATC CTT CCT CGT CCC | 1723 |
| PSD2 KO check R | GTCTCCGAACGAATTGAGAATCTG | 4758 |
| PSD2 KO F | GTA AAG AAT CCT CGA TTT TCA GGA GCA TCC AAC GAC GAA GAG ATT GTA CTG AGA GTG CAC | 1721 |
| PSD2 KO R | ATT TTG GTA ACC ACT AAC TAC AGC CAA TTT TTC GGC GGC TCT GTG CGG TAT TTC ACA CCG | 1722 |
| TAZ1 KO check F | CTGGCCAGTTAGAAGGTACAGCATAATCAA | 5030 |
| TAZ1 KO check R | CACTCGTATGGCACACTAAATC | 5031 |
| TAZ1 KO F | TCATTTTCAAAAAAAAAAAAAGTAAAGTTTTCCCTA TCAAagattgtactgagagtgcac | 5028 |
| TAZ1 KO R | CATACATGCTAGTATTTACACGAATTTAATTGCTTA AATTctgtgcggtatttcacaccg | 5029 |
| Tim50 Tag F | TGAAGAGGAAAAGAAAAAGAAGAAGATTGCTGAAT CCAAAGGTGACGGTGCTGGTTTA | 419 |
| Tim50 Tag R | ATAGATACGTAGATACATGAGAAGAGGGTTTACAT GAAAATCGATGAATTCGAGCTCG | 420 |

| | | |
|---|---|---|
| Tom70 Tag F | TCAAGAAACTTTAGCTAAATTACGCGAACAGGGTT TAATGGGTGACGGTGCTGGTTTA | 797 |
| Tom70 Tag R | TTTGTCTTCTCCTAAAAGTTTTTAAGTTTATGTTTAC TGTTCGATGAATTCGAGCTCG | 798 |
| UPS1 KO check F | AACCGGAATCAAGCACCAAGGTAGTAAGCA | 745 |
| UPS1 KO check R | CTTGGATGTTCTTCGGCAACAC | 4757 |
| UPS1 KO F | TGGCTTCTGAGACGGCGGTAAGATATCCTTAAGAG TTGCAagattgtactgagagtgcac | 743 |
| UPS1 KO R | CGCCCATGGTGATATCTTTAAAGATCTTTAAATGGG AACActgtgcggtatttcacaccg | 744 |
| UPS2 KO check F | ATCTTGAATGAGATGATTATGTGCGATGGC | 4478 |
| UPS2 KO check R | GTGGTAAGTGTCATTCCGTCTAC | 4479 |
| UPS2 KO F | AGACTAAGATAAAATAATCGAGAATAATTAAAAGAC GATAagattgtactgagagtgcac | 4476 |
| UPS2 KO R | GTAGTATGCAGTGCCATGCGGGATCAAGGAATTTG TATCTctgtgcggtatttcacaccg | 4477 |
